# YIPPEE/YPEL5 regulates circadian period by modulating PER ubiquitination and stability

**DOI:** 10.64898/2026.04.28.721296

**Authors:** Meilin Li, Yue Zhou, Huihui Huang, Mengmeng Ren, Wen Hu, Xianhui Liu, Yong Zhang

## Abstract

Circadian clocks rely on evolutionarily conserved transcriptional–translational feedback loops to align physiology and behavior with the 24-hour day–night cycle. Post-translational modifications, especially ubiquitination, critically control core clock protein stability and circadian pace. In both fruit flies and mammals, the abundance of key pacemaker protein PERIOD (PER/PER2) must be tightly regulated to sustain circadian rhythms. However, the canonical phosphorylation-dependent pathway cannot fully account for turnover of PER in *Drosophila*. Here we identify Yippee as a critical regulator of PER level through a genetic screen of downregulating ubiquitination pathways. Knockdown of *Yippee* in circadian pacemaker neurons significantly lengthens the circadian period by ∼2 hours and elevates PER levels. Indeed, YIPPEE physically interacts with PER in fly head. Mechanistically, its mammalian homolog YPEL5 binds PER2, promotes K63-linked polyubiquitination and proteasomal degradation of PER2 independently of FASP-region phosphorylation. Lastly, downregulating *YPEL5/Ypel5* prolongs circadian periods in human U2OS cells and mice. Our findings reveal an evolutionarily conserved mechanism in which Yippee/YPEL5 regulates circadian rhythms by mediating ubiquitin-dependent degradation of PER/PER2.

## Introduction

To adapt to the predictable environmental changes caused by Earth’s rotation, most organisms have evolved an endogenous circadian timing system, which operates on a ∼24-hour cycle. The circadian clocks help organisms adjust to daily environmental fluctuations by synchronizing daily rhythms of physiology and behavior^[1]^. In mammals, circadian rhythms are coordinated by the central pacemaker located in the suprachiasmatic nucleus (SCN) of the hypothalamus^[2]^. In *Drosophila melanogaster*, circadian rhythms are primarily regulated by about 150 clock neurons located in the fly brain^[3]^. Among these neurons, the small ventrolateral neurons (sLNvs) are termed pacemaker neurons, as they synchronize other clock neurons by secreting the pigment-dispersing factor (PDF)^[4, 5]^. At the molecular level, circadian clock is maintained by a highly conserved transcription-translation feedback loop (TTFL) across species^[6]^. In *Drosophila*, the transcriptional activators, CLOCK (CLK) and CYCLE (CYC), promote the expression of PERIOD (PER) and TIMELESS (TIM), which inhibit CLK/CYC activity after dimerization^[7, 8]^. A similar TTFL operates in mammals, in which CLOCK and BMAL1 activate the expression of *Per* and *Cry* genes, and the resulting PER/CRY complexes subsequently repress their own transcription^[9, 10]^.

As the principal transcriptional repressors in the TTFL, PER in flies and PER2 in mammals are central determinants of circadian speed and robustness^[11, 12]^. In particular, PER protein levels are tightly correlated with the speed of molecular oscillators. Post-translational modifications (PTMs) of PER have been shown to play fundamental roles in regulating the pace of circadian clocks, such as glycosylation and phosphorylation^[13, 14]^. In both flies and mammals, casein kinase 1 (CK1) family kinases (*Drosophila* homolog DOUBLETIME, DBT) phosphorylate PER proteins, thereby regulating their stability, subcellular localization, and repressor activity, making phosphorylation a central mechanism for period determination^[15, 16]^.

PER is degraded by the ubiquitin ligase complex containing the F-box protein SLIMB (the *Drosophila* homolog of mammalian β-TrCP), via phosphorylation-dependent ubiquitination^[17–19]^. However, PER abundance still exhibits daily oscillation even after SLIMB loss, albeit with a compromised amplitude compared to wild-type flies^[18]^. Therefore, it is likely that additional ubiquitin-related factors besides SLIMB/β-TrCP regulate PER protein degradation and remain to be discovered.

To systematically characterize ubiquitin-mediated regulation of circadian rhythms, we performed a genetic screen using *Drosophila* lines with perturbations in ubiquitination-related genes. We found that knockdown of *Yippee* in clock neurons significantly lengthened the circadian period.

YIPPEE is highly conserved between *Drosophila* and mammals (70.8% amino acid sequence identity) and participates in processes like cell proliferation, immune response and tumorigenesis, but its molecular function is poorly understood^[20–23]^. A drug target study suggested that thalidomide inhibits a ubiquitin ligase complex by targeting cereblon’s CULT domain, which is highly homologous to *Drosophila* YIPPEE^[24]^. This observation has led to the hypothesis that *Drosophila* YIPPEE may function as a component of ubiquitin ligase complex, although direct evidence remains lacking.

Here we demonstrate that YIPPEE plays an essential role in regulation of circadian rhythms. Knockdown of *Yippee* in clock neurons resulted in a significantly prolonged behavioral period in *Drosophila* with elevated PER abundance. Indeed, YIPPEE and its mammalian homolog YPEL5 physically interact with dPER/hPER2 and repress their protein stability. Furthermore, our results indicate that YIPPEE/YPEL5 promotes the ubiquitin-mediated degradation of dPER/hPER2. Subsequent experiments confirmed that YIPPEE/YPEL5 promote K63-linked polyubiquitination of dPER/hPER2 independently of phosphorylation within the PER2 FASP region. Finally, specific knockdown *Ypel5* in mouse SCN and *YPEL5* in human U2OS cells demonstrates that *Yippee*-mediated circadian regulation is conserved across species. Together, these findings establish YIPPEE/YPEL5 as an evolutionarily conserved regulator of PER turnover and circadian timing.

## Results

### *Yippee* downregulation lengthens circadian locomotor period

In order to identify novel regulator of *Drosophila* circadian rhythms, we conducted a RNAi screen of 56 lines targeting ubiquitin-related genes in clock neurons. This screen identified two candidates that markedly extended the circadian locomotor period: *Ctrip* (circadian trip) and *Yippee*, with both producing the longest period (Figure 1A). Since *Ctrip* has previously been implicated in CLOCK stability^[25]^, this finding validates the reliability of our screen. We therefore focused on investigating the role of *Yippee* in circadian regulation in the current study.

**Figure 1.**
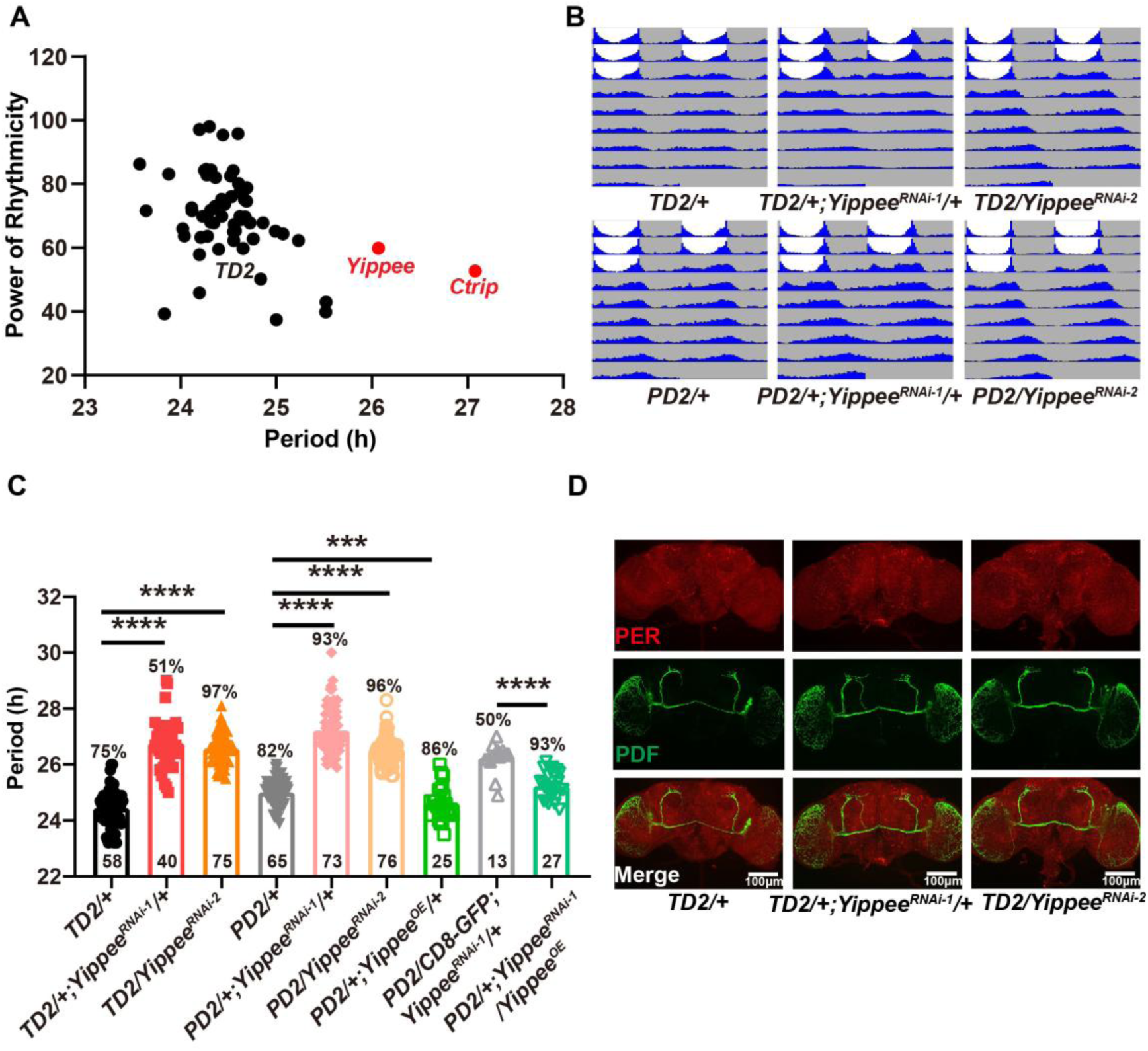
Downregulation of *Yippee* in the circadian clock leads to a long-period phenotype. (A) A screen of *Drosophila* ubiquitin-related genes identified *Yippee* as a candidate gene. *TD2* denotes *tim-GAL4, UAS-Dicer2*; *Yippee* denotes TD2>UAS-*Yippee* RNAi flies; *Ctrip* denotes TD2>UAS-*Ctrip* RNAi flies. (B) Representative actograms showing lengthened circadian periods in *Yippee* knockdown flies. *Yippee* was downregulated in all circadian neurons (top) and specifically in PDF^-^ positive neurons (bottom). *PD2* denotes *Pdf-GAL4, UAS-Dicer2*. (C) Quantification of behavioral circadian period following *Yippee* knockdown or overexpression in clock neurons. *Yippee^OE^* denotes *UAS-Yippee-3×HA; CD8-GFP* denotes *UAS-mCD8::GFP.L.* Numbers within bars indicate the number of rhythmic flies, while percentages indicate the proportion of rhythmic individuals among total tested. Statistical analysis was performed using one-way ANOVA. Error bars represent SEM. ***: *P* < 0.001, ****: *P* < 0.0001.(D) Representative confocal images of *Yippee* RNAi flies. Flies were entrained under three light–dark (LD) cycles, dissected at Zeitgeber Time 1 (ZT1, ZT0 represents when light is turned on, while ZT12 indicates when light is turned off), and immunostained with anti-PDF (green) and anti-PER (red) antibodies.

To mitigate the potential off-target effects of RNAi, we included two independent lines targeting different exons of the *Yippee* gene, both of which exhibited significant knockdown efficiency when driven by a pan-neuronal driver (Figure S1A-B). Indeed, knockdown *Yippee* in all clock neurons (*TD2>UAS-Yippee RNAi*) resulted in an approximately 2-hour extension of the free-running period with both RNAi lines (Figure 1B-C). To identify the neuronal clusters in which *Yippee* functions, we further downregulated *Yippee* in two distinct clock neuron populations: PDF^+^ neurons (*PD2>UAS-Yippee RNAi*), or PDF^−^ neurons (*TD2; Pdf-GAL80>UAS-Yippee RNAi*). Notably, specific knockdown of *Yippee* in PDF^+^ neurons yielded a similar period lengthening, whereas knockdown in PDF^−^ neurons only induced a modest extension of ∼0.5 hour (Figure 1B–C, S1C). These results suggest that *Yippee* primarily acts in PDF^+^ circadian neurons to regulate locomotor period.

To further validate the effect of *Yippee* on circadian clock, we utilized the *Yippee^OE^-3×HA* transgene to overexpress *Yippee* (Figure S1D). Interestingly, overexpression of *Yippee* in PDF^+^ neurons resulted in a modest but significant shortening of the circadian period compared to control (Figure 1C). Furthermore, co-expression of the *Yippee* transgene rescued the long-period defects induced by *Yippee* knockdown, which reinforces the functional link between *Yippee* expression levels and circadian period length (Figure 1C).

Lastly, to exclude the possibility that *Yippee* knockdown causes developmental defects in PDF^+^ circadian neurons, we examined the axonal projections of these neurons. We observed no apparent abnormalities in the major projections of PDF neurons (Figure 1D), suggesting that the behavioral phenotypes are not attributed to gross structural defects of PDF^+^ neurons. Together, these findings indicate that *Yippee* regulates the circadian period primarily through PDF^+^ neurons, prompting us to explore its potential influence on core circadian clock components at the molecular level.

### PER abundance is increased in *Yippee* knockdown and contributes to the period lengthening

Since *Yippee* knockdown led to a long-period phenotype, we hypothesized that it might alter the abundance of core clock genes. We firstly analyzed the expression pattern of *per*, *tim*, and *clk* in *Yippee* knockdown flies using RT-qPCR at different time of the day. Our results showed that the circadian oscillation and abundance of these pacemaker genes were not significantly changed (Figure S2A–C). Therefore, we speculated that *Yippee* may affect the molecular clock at the post-translational level.

Next, we performed immunofluorescence staining with anti-PER antibodies in sLNvs at Zeitgeber time 1 (ZT1). Indeed, both *Yippee* RNAi lines exhibited a significant increase in PER levels by approximately 2-3 folds (Figure 2A). To further determine whether *Yippee* knockdown affects overall PER oscillation and to rule out potential effects of light on PER levels, we placed flies in constant darkness for three days and performed immunostaining of fly brains every four hours. Consistent with the observation at ZT1, PER levels remained elevated throughout the circadian cycle compared to controls, and the phase of PER oscillation was delayed by more than four hours (Figure 2B), consistent with the long-period behavioral phenotype.

**Figure 2.**
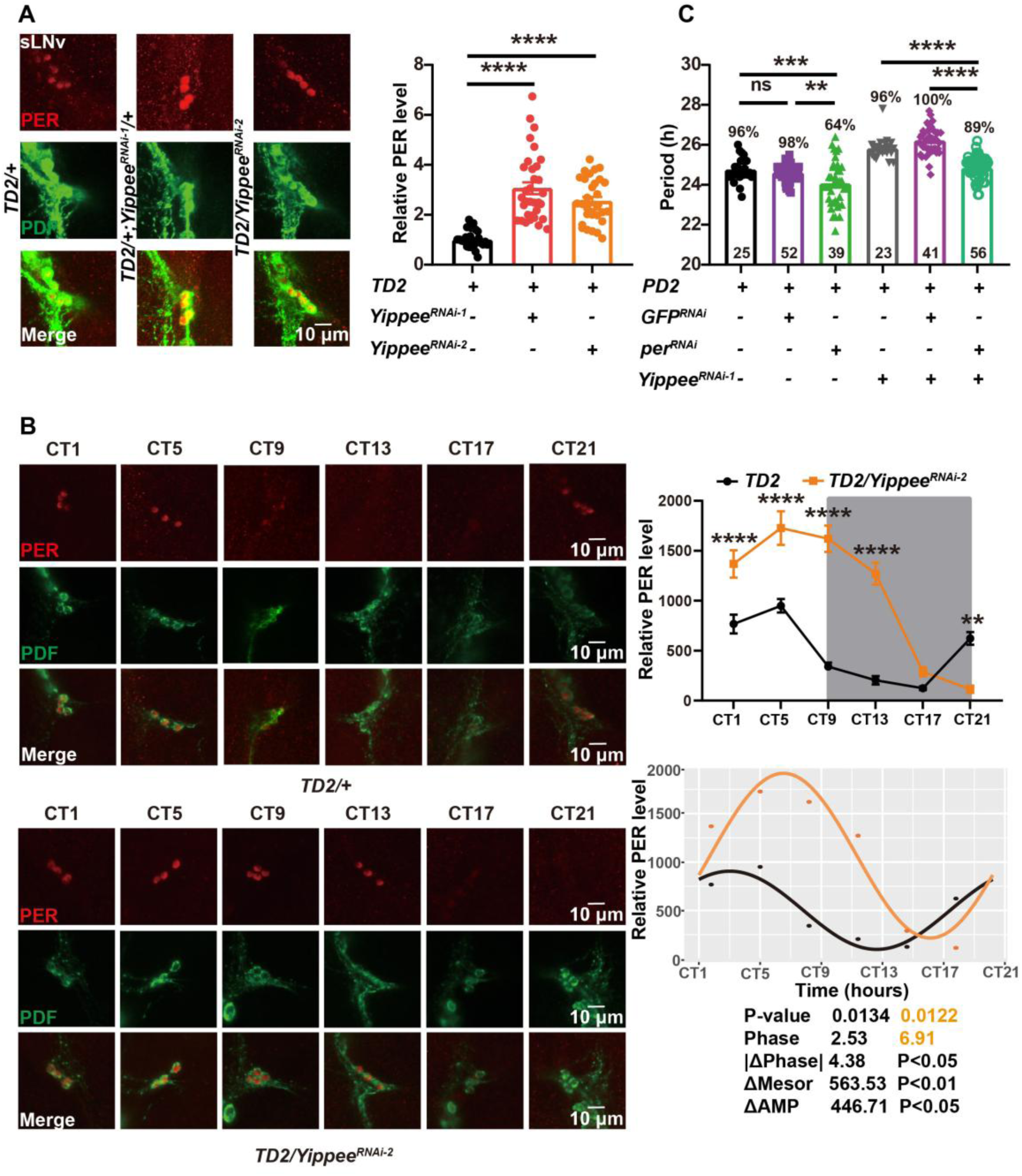
*Yippee* regulates circadian period by increasing PER protein levels. (A) Representative confocal images showing PER and PDF expression in the small ventrolateral neurons (sLNvs). Flies were entrained under three light–dark (LD) cycles, adult brains were dissected at ZT1, and PDF staining was used to locate sLNvs. Relative PER levels were quantified (right panel). Statistical analysis was performed using one-way ANOVA. Error bars represent SEM. ****: *P* < 0.0001. (B) Representative images of PER protein oscillation in sLNvs. *TD2/+* and *TD2/Yippee^RNAi-2^* flies were entrained under light–dark (LD) conditions for three days, followed by constant darkness (DD) for four days. Sampling began at Circadian Time 1 (CT1) and continued every four hours. The top-right panel shows quantitative analysis. Statistical analysis was performed using two-way ANOVA. Error bars represent SEM. **: *P* < 0.01, ****: *P* < 0.0001. The bottom-right panel was generated using R Studio and the oscillating curves were compared using with Circacompare to gain the difference of peak time (|Δphase|), expression (Δmesor) and amplitude (ΔAMP). (C) Graph showing the circadian period of *Yippee* and *per* double knockdown flies. *GFP^RNAi^*denotes *UAS-GFP* RNAi, and *per^RNAi^* denotes *UAS-per* RNAi. Statistical analysis was performed using one-way ANOVA and unpaired *t*-tests. Error bars represent SEM. **: *P* < 0.01, ***: *P* < 0.001, ****: *P* < 0.0001; ns, not significant.

To further test the hypothesis that *Yippee* knockdown lengthens the circadian period by elevating PER protein levels, we generated flies with simultaneous knockdown of *Yippee* and *per* (using GFP RNAi as a control), and monitored their locomotor rhythms. Interestingly, the long-period phenotype was rescued in these double knockdown flies (Figure 2C). Consistently, immunostaining showed that the elevated PER levels in *Yippee* knockdown flies were reduced by co-silencing *per*, supporting the interpretation that the rescue effect was accompanied by *per* downregulation (Figure S3A-B). Together, these findings indicate that *Yippee* modulates the circadian period by decreasing the protein abundance of PER.

### YIPPEE physically interacts with PER and may be involved in the ubiquitin machinery

Next, we sought to investigate the potential molecular mechanism by which YIPPEE regulates PER abundance. To this end, we overexpressed the *Yippee^OE^-3×HA* in circadian neurons (*TD2/+; Yippee^OE^-3×HA/+*) and performed immunoprecipitation followed by mass spectrometry (IP-MS) using anti-HA antibodies on fly head extracts to identify potential proteins interacting with YIPPEE.

Consistent with the predicated function of YIPPEE in a ubiquitin ligase complex, Gene Ontology (GO) enrichment analysis supported YIPPEE’s involvement in proteasome-related processes (Figure 3A). STRING (https://string-db.org/) analysis of YIPPEE interactome further revealed that UbcE2H–a predicted ubiquitin-conjugating enzyme (E2), and CG31357, an E3 ubiquitin ligase, interact with YIPPEE (Figure 3B). These results suggest YIPPEE may be part of the ubiquitin machinery. However, PER was not detected among the IP-MS hits, likely due to the transient or weak nature of protein–protein interactions or low abundance of this protein. We therefore performed co-immunoprecipitation (Co-IP) using head extracts from flies overexpressing both *Yippee* and *per* (*TD2/+; Yippee^OE^-3×HA/per^24^*). Indeed, YIPPEE physically interacts with PER (Figure 3C). Taken together, combined with our observation that *Yippee* knockdown elevates PER levels, we speculated that YIPPEE might physically associate with PER to promote its proteasome-mediated degradation.

**Figure 3.**
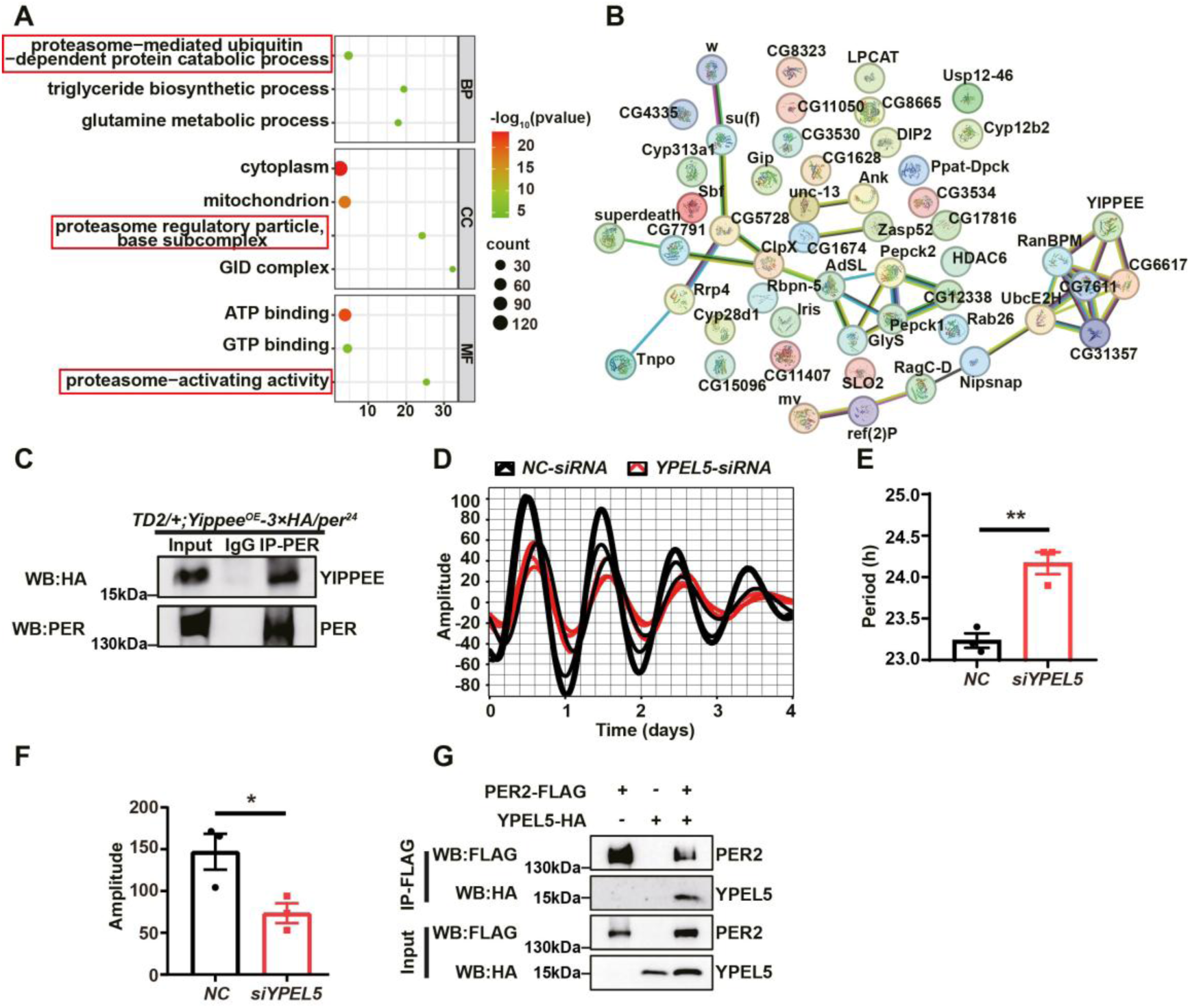
YIPPEE associates with PER and the regulatory role of *Yippee* in PER protein is conserved in mammalian cells. (A) Gene Ontology (GO) enrichment analysis of YIPPEE-interacting proteins identified by immunoprecipitation–mass spectrometry (IP-MS). Displayed GO terms met the significance thresholds of *P* < 0.05 and false discovery rate (FDR) < 0.05. GO categories include Biological Process (BP), Cellular Component (CC), and Molecular Function (MF). (B) Network analysis of YIPPEE-interacting proteins using STRING analysis. Proteins were selected based on a mean enrichment ratio ≥ 2 and *P* < 0.05. (C) Co-immunoprecipitation (Co-IP) showing interaction between YIPPEE and PER in *TD2/+*; *Yippee^OE^-3×HA/per^24^* fly heads collected at ZT1. “Input” indicates total lysate; “IgG” group was immunoprecipitated with anti-IgG; “IP” group was immunoprecipitated with anti-PER antibody. (D-F) Graph showing the oscillation of PER2::LUC signals in human U2OS cells transfected with either negative control (*NC*) or *siYpel5*. Statistical analysis was performed using unpaired *t*-tests. Error bars represent SEM. *: *P* < 0.05, **: *P* < 0.01. (G) Co-IP analysis showing interaction between YPEL5 and PER2 in HEK293T cells. “Input” indicates total cell lysate; “IP” group used anti-FLAG for immunoprecipitation.

### The regulation of *Yippee* on circadian rhythm and PER is conserved among species

Given *Yippee*’s role in regulating PER and circadian rhythm in *Drosophila*, we next examined whether its circadian function is conserved in mammals. The YIPPEE protein family is evolutionarily conserved^[26]^, and our sequence alignment confirmed this conservation across flies, mammals and even plants (Figure S4A). We first utilized the *PER2*-luciferase U2OS cell line to monitor circadian oscillations under *YPEL5* (the human homolog of *Drosophila* YIPPEE) knockdown conditions. Interestingly, consistent with the lengthened behavioral period observed in *Yippee* knockdown flies, downregulation of *YPEL5* in U2OS cells (*siYPEL5*) resulted in a significantly lengthened period (approximately one hour) compared to the control siRNA (*NC*) (Figure 3D-E). We also observed a marked reduction in the amplitude of circadian oscillation, which decreased to nearly half of the control (Figure 3D, F).

We next tested whether the human YPEL5 could also interact with PER2. After confirming the expression of PER2-FLAG and YPEL5-HA in HEK293T cells (Figure S4B–C), we co-transfected these two constructs and conducted Co-IP assays. Indeed, we detected a physical interaction between YPEL5 and PER2 (Figure 3G), suggesting that the regulatory relationship between YIPPEE and PER is likely conserved across species.

### YPEL5 regulates PER2 stability via K63-linked polyubiquitination in mammalian cells

Inspired by these findings, we examined whether YPEL5 modulates PER2 stability in mammalian cells. In HEK293T cells, cycloheximide (CHX) treatment to inhibit protein synthesis led to gradual decreases in PER2 protein levels over time. However, *YPEL5* knockdown noticeably slowed PER2 degradation (Figure 4A-B), mirroring the PER increase observed in *Yippee* knockdown flies.

**Figure 4.**
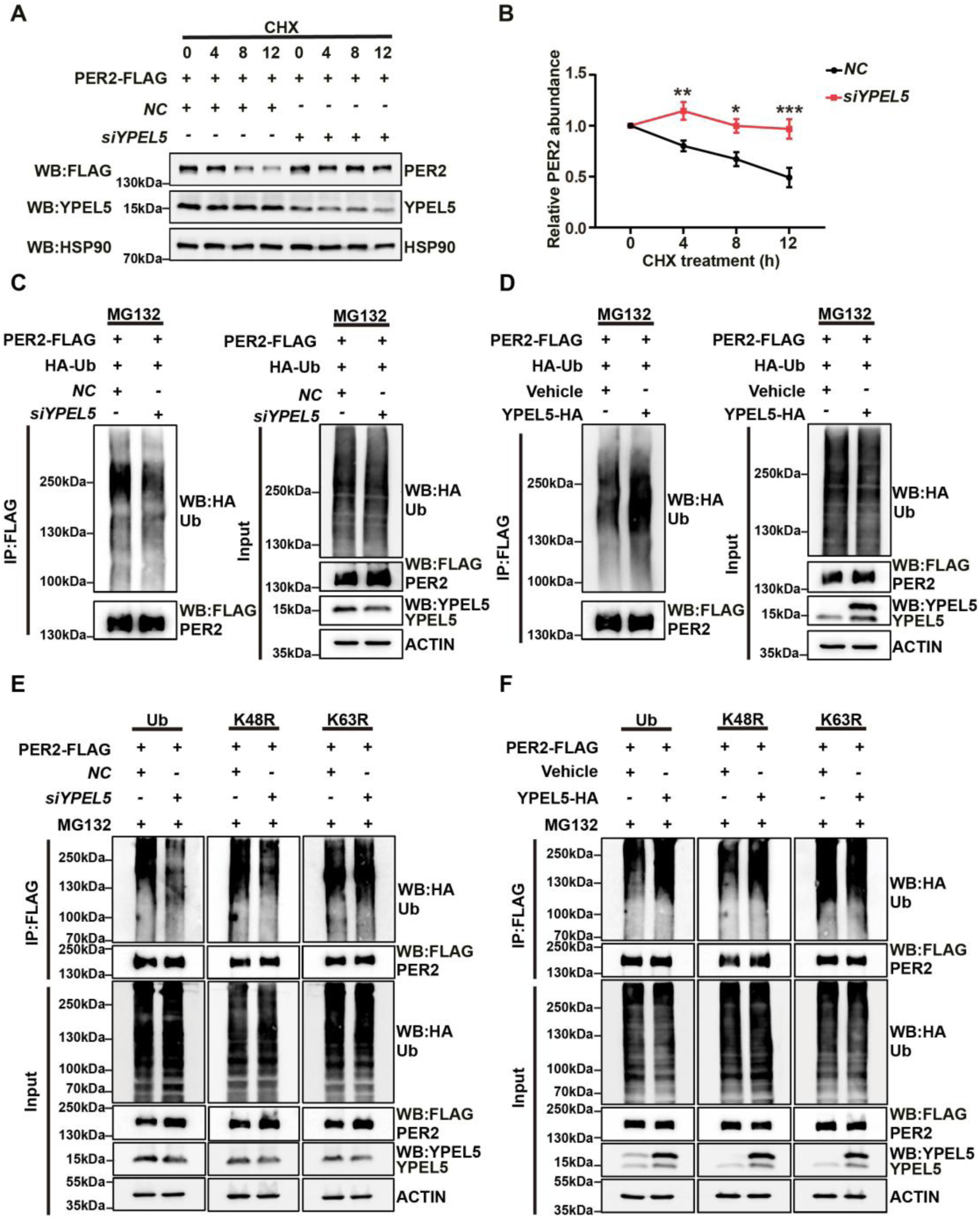
YPEL5 promotes PER2 ubiquitination through K63-linked polyubiquitin modification. (A-B) Cycloheximide chase assay to illustrate the degradation rate of PER2 in HEK293T cells transfected with or without *siYPEL5.* Cells were treated with 100 μg/mL CHX. HSP90 was used as a loading control. The quantification is shown on the right. Statistical analysis was performed using two-way ANOVA followed by Dunnett’s multiple comparisons test. Error bars represent SEM. *: *P* < 0.05, **: *P* < 0.01, ***: *P* < 0.001. (C-D) Representative blots showing the ubiquitination of PER2 in HEK293T cells with *Ypel5* knockdown (C) or overexpression (D). The left panels show the results from immunoprecipitated PER2, while the right panels show total protein levels in cell lysates. (E-F) Representative blots showing the ubiquitination of PER2 in HEK293T cells transfected with wild-type Ub, K48R, or K63R ubiquitin mutants. *YPEL5* is either knocked down (E) or overexpressed (F) to show the effect of YPEL5 on different types of PER2 ubiquitination.

Given YIPPEE/YPEL5’s close association with the ubiquitin–proteasome pathway, we next determined whether YPEL5 regulates PER2 ubiquitination. Using MG132 to inhibit proteasomal degradation and allow accumulation of ubiquitinated PER2, we performed ubiquitination assays under *YPEL5* knockdown and overexpression conditions. Immunoprecipitation of PER2 followed by ubiquitin immunoblotting revealed that *YPEL5* depletion decreased whereas *YPEL5* overexpression increased PER2 ubiquitination (Figure 4C–D, replicates in Figure S5A–B). These data suggest that YPEL5 promotes PER2 degradation by enhancing its ubiquitination.

Ubiquitin conjugation via different lysine residues forms polyubiquitin chains with distinct cellular fates. K48-linked polyubiquitinchains typically mediate proteasomal degradation, while K63-linked polyubiquitinchains are often associated with non-degradative functions such as signaling or trafficking.^[27, 28]^ To identify the linkage type involved in YPEL5-mediated PER2 regulation, we performed ubiquitination assays using wild-type ubiquitin and the lysine-to-arginine mutants K48R and K63R in cells co-transfected with PER2 and control or YPEL5 constructs. With wild-type ubiquitin, *YPEL5* knockdown reduced PER2 ubiquitination (Figure 4E), validating our prior findings. Notably, *YPEL5* knockdown still led to a decrease in PER2 ubiquitination in the K48R group, indicating that K48 is non-essential. In contrast, disrupting K63-linked ubiquitination abrogated the effect of *YPEL5* knockdown on PER2 ubiquitination (Figure 4E). Similarly, *YPEL5* overexpression increased PER2 ubiquitination with wild-type and K48R ubiquitin, but not with the K63R mutant (Figure 4F). Independent replicates confirmed these observations (Figure S5C–D), demonstrating YPEL5 specifically promotes K63-linked PER2 ubiquitination, suggesting a non-canonical role for K63-linked polyubiquitination in targeting PER2 for proteasomal degradation.

### YPEL5–PER2 interaction is independent of phosphorylation within the PER2 FASP region

PER2 degradation follows a biphasic circadian pattern: newly synthesized PER2 is degraded in the cytoplasm to avoid premature inhibition of BMAL1/CLOCK, whereas nuclear PER2 is modified and targeted for degradation late in the daily cycle to relieve transcriptional repression^[29, 30]^. We thus mapped YPEL5–PER2 interaction by subcellular fractionation and Co-IP, which showed their association in both the cytoplasm and nucleus (Figure S6A–B), indicating the YPEL5 regulates PER2 in both compartments.

To further map the region of PER2 responsible for YPEL5 binding, we generated a series of PER2 truncated constructs covering the N-terminus (1–500 aa), central region (501–950 aa), and C-terminus (951–1255 aa) fragments. The central fragment contains the well-characterized CK1ε/δ phosphorylation cluster, Ser662–Ser674, and the expression of all truncated variants was verified (Figure 5A, Figure S6C). Co-IP assays demonstrated that YPEL5 interacted with both the N-terminal and C-terminal regions of PER2, whereas no binding was detected the central 501–950 aa fragment (Figure 5B). Phosphorylation plays a critical role in regulating PER2 ubiquitination by specific E3 ligases. Mechanistically, CK1ε/δ-dependent phosphorylation at PER2 Ser480 facilitates β-TrCP recognition, and subsequent ubiquitination, while phosphorylation at Ser662 and adjacent residues suppresses β-TrCP dependent degradation^[19, 29, 31]^. These distinct regulatory patterns indicate that PER2 phosphorylation exerts divergent effects on E3 ligase recruitment. Given that the 501–950 aa region harbors multiple phosphorylation sites, including the familial advanced sleep phase (FASP) region (S662–674), we next explored whether phosphorylation within this region governs YPEL5-PER2 interaction. We generated phospho-deficient (PER2 Ser662-674A) and phospho-mimetic PER2 (S662–674D) mutants (Figure S6D) and performed Co-IP analysis. Functional validation revealed that both phospho-deficient and phospho-mimetic mutants retained robust binding capacity to YPEL5 (Figure 5C). Collectively, these results indicate that the YPEL5–PER2 interaction is independent of phosphorylation within the PER2 FASP region.

**Figure 5.**
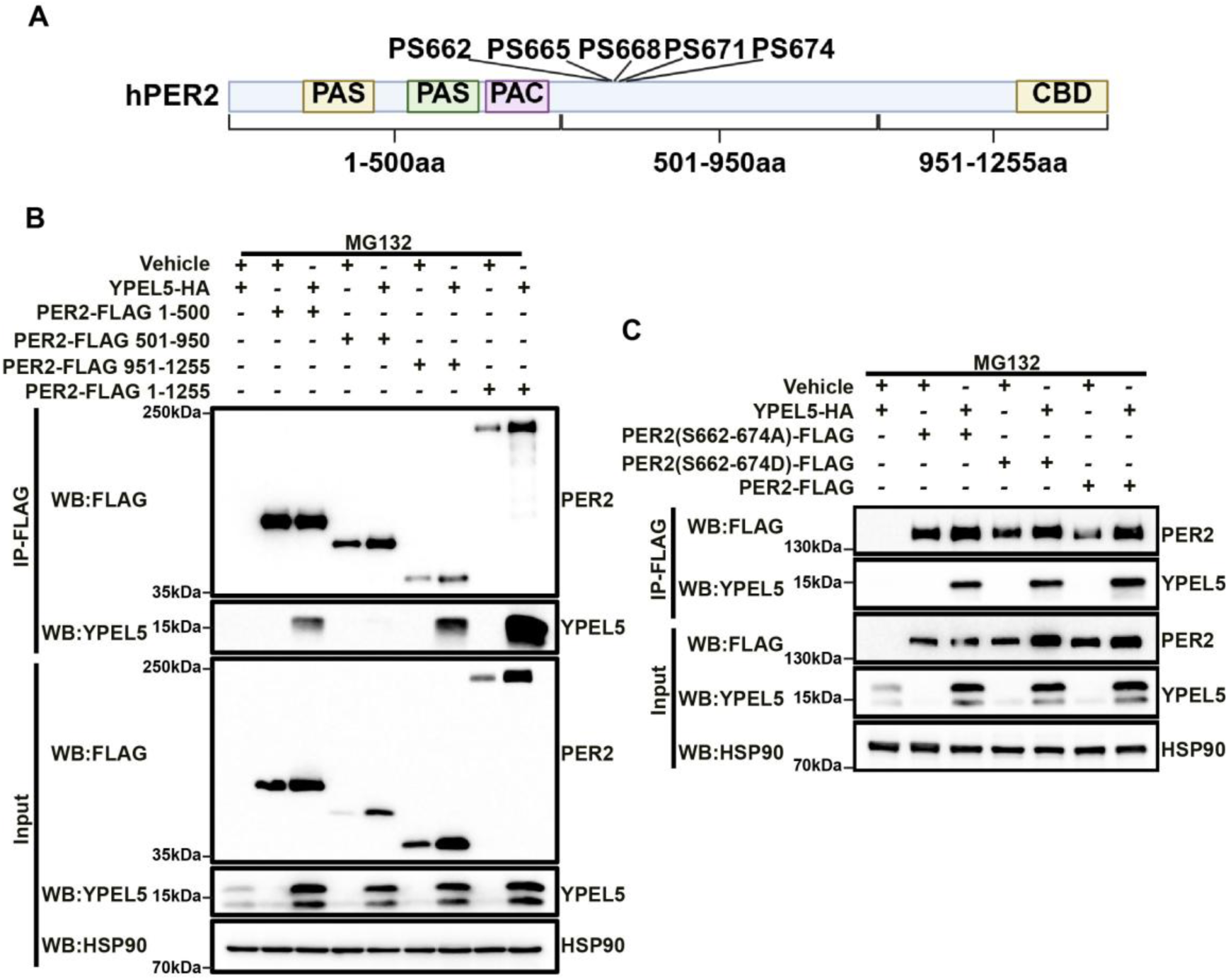
Binding of PER2 by YPEL5 is independent of phosphorylation within the PER2 FASP region. (A) Schematic illustration of the PER2 constructs used in panel (B) and (C). The fragments 1–500 aa, 501–950 aa, and 951–1255 aa represent distinct regions of PER2. S662, S665, S668, S671, and S674 are the reported phosphorylation residues within PER2. (B) Analysis of the interaction between YPEL5 and different structural domains of PER2. The top panels show the results from immunoprecipitated PER2, while the bottom panels show total protein levels in cell lysates. (C) Representative blots showing the interaction between YPEL5 and PER2(S662–674A) or PER2(S662–674D). PER2(S662–674A) denotes alanine substitution of residues S662, S665, S668, S671, and S674, whereas PER2(S662–674D) denotes aspartic acid substitution at these sites.

### *Yippee* regulates activity rhythms in mice

Given that our *Drosophila* and cellular data indicate YIPPEE/YPEL5 regulates circadian rhythms and PER protein stability, we next examined whether YPEL5 modulates mouse circadian locomotor activity. To achieve tissue-specific knockdown of *Ypel5* in mice, we utilized *Ypel5^Flox/Flox^*homozygous mice and bilaterally injected VGAT1-eGFP (control) or VGAT1-Cre viruses into their suprachiasmatic nucleus (SCN)—the master circadian pacemaker in the mouse brain (Figure 6A). Immunofluorescence staining verified viral targeting to the SCN (Figure 6B). Following three weeks of viral expression, we monitored the locomotor rhythms using a running wheel assay under constant darkness. Compared to the control, VGAT1-Cre–injected mice exhibited delayed activity onset, more scattered patterns of behavior, and even bouts of activity during the subjective day (Figure 6B). Quantification of their circadian period length revealed a 0.7-hour lengthening in *Ypel5* knockdown mice (Figure 6C). Together, these findings demonstrate that *YPEL5*/*Ypel5* depletion prolongs circadian period in *Drosophila*, human cells, and mice, supporting an evolutionarily conserved function across species.

**Figure 6.**
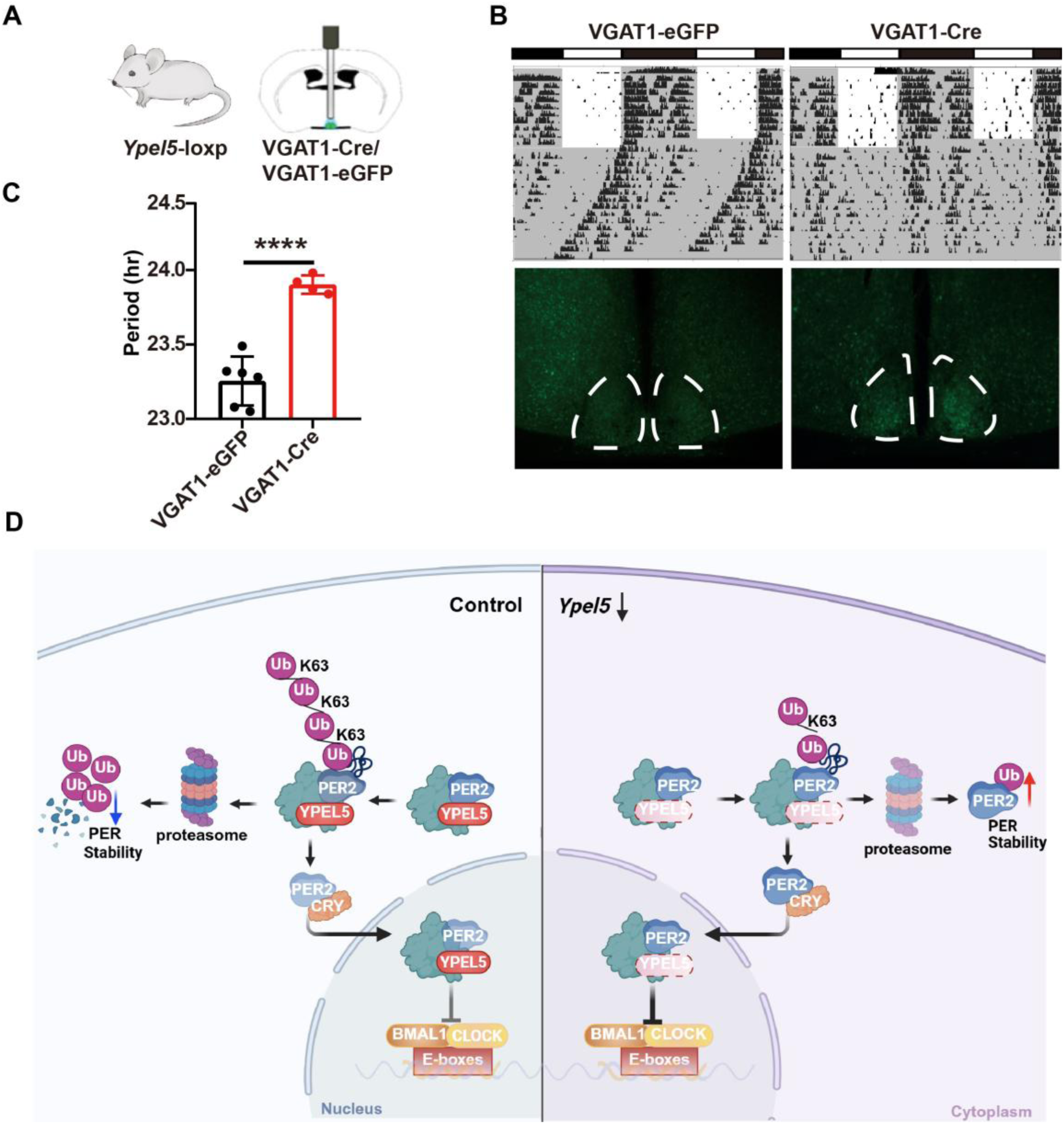
The role of *Yippee* in circadian regulation is conserved across species. (A) Schematic illustration of site-specific brain injection targeting the SCN in mice. *Ypel5^Flox/Flox^* homozygous mice received stereotaxic injections into the suprachiasmatic nucleus (SCN). Control mice were injected with VGAT1-eGFP virus, and experimental mice received VGAT1-Cre virus to induce conditional knockout. (B) Double plotted actogram showing the locomotor activity rhythm of mice injected with either VGAT1-eGFP or VGAT1-Cre. (C) Quantification of the circadian periods in panel (B). Statistical analysis was performed using unpaired t-tests. Error bars represent SEM. ****: *P* < 0.0001. (D) Proposed working model of *Yippee* function in circadian regulation. YIPPEE binds to PER and promotes its K63-linked ubiquitination and subsequent degradation.

## Discussion

In this study, we conducted a genetic screen and identified YIPPEE as a previously uncharacterized post-translational regulator of circadian rhythms. Behaviorally, depleting *Yippee* expression in *Drosophila* circadian neurons and mouse SCN markedly lengthened the circadian period. Mechanistically, YPEL5, the human ortholog of YIPPEE, binds to the core clock protein PER2 and promotes its K63-linked polyubiquitination, thereby modulating PER2 stability. Together, these findings reveal an evolutionarily conserved pathway whereby YIPPEE/YPEL5 fine-tunes circadian rhythms by governing PER/PER2 protein turnover. Specifically, YIPPEE/YPEL5 interacts with PER/PER2 to facilitate its ubiquitination and subsequent degradation, ultimately shaping circadian period length (Fig. 6D).

We identified YIPPEE as a novel regulator that reduces PER protein stability. To date, SLIMB is the most well-characterized E3 ligase responsible for the degradation of PER in *Drosophila*^[19, 32]^. Notably, loss of SLIMB fails to fully block PER turnover^[18]^, implying the existence of additional regulatory factors. Early sequence analyses predicted that YIPPEE may function within a ubiquitin ligase complex. Here, we experimentally validated YIPPEE as a critical modulator of protein stability. Although YIPPEE lacks canonical E3 ligase domains, it likely acts as an adaptor protein. Our IP-MS profiling identified multiple E3 ligases that interact with YIPPEE, such as CG31357. Whether these E3 ligases mediate YIPPEE-dependent PER regulation remains to be further investigated.

One of the most unexpected findings of this study is that YPEL5 promotes K63-linked polyubiquitination of PER2. Traditionally, K63-linked polyubiquitin chains are associated with non-degradative processes, such as signal transduction and intracellular trafficking, whereas K48-linked polyubiquitin chains serve as the canonical signal for proteasomal degradation^[33]^. However, emerging evidence suggests that K63-linked ubiquitination can also participate in protein degradation under certain contexts. For example, K63 ubiquitination has been shown to facilitate proteasome-dependent degradation by serving as a seed for the assembly of K48/K63-branched ubiquitin chains, and it can also act as a signal for chaperone-mediated autophagic degradation of specific substrates^[34, 35]^. In this regard, our results extend this concept to the circadian system and suggest that K63-linked polyubiquitination of PER2 is not merely a non-degradative modification. Instead, it may represent an intermediate or instructive signal that promotes PER2 turnover. One possibility is that YPEL5-mediated K63 polyubiquitin chains on PER2 facilitate the recruitment of additional ubiquitination machinery, thereby enabling the formation of branched ubiquitin chains that direct PER2 to the proteasome. Although the precise downstream fate of K63-ubiquitinated PER2 remains to be determined, our findings broaden the current understanding of how distinct ubiquitin linkages regulate clock protein stability and highlight a previously unappreciated layer of post-translational control in circadian timing.

Our cross-species data demonstrates that *Ypel5* knockdown in the mouse SCN disrupts circadian locomotor activity, while human YPEL5 interacts with PER2 to regulate its stability and oscillation in cultured cells. This evolutionary conservation underscores the importance of the YIPPEE–PER regulatory axis in circadian biology and raises intriguing questions about its potential relevance to human health. Given that both circadian dysregulation and aberrant YIPPEE/YPEL5 expression are linked to oncogenesis and immune regulation, it will be valuable to explore whether YPEL5 mediates the connection between circadian rhythm disruption and pathological conditions.

In summary, our results strongly support a novel role for YIPPEE/YPEL5 in promoting PER/PER2 degradation to regulate circadian period length. However, several key questions remain unanswered. First, how does YPEL5 enhance PER2 degradation by promoting K63-linked ubiquitination—a modification traditionally associated with signal transduction rather than proteasomal degradation? Second, under which (patho)physiological contexts do YIPPEE/YPEL5 and SLIMB/β-TrCP differentially regulate PER degradation? Finally, it will be critical to determine whether YIPPEE/YPEL5 targets additional protein substrates beyond PER and the circadian clock, as well as the mechanism by which YIPPEE/YPEL5 modulates cellular biology.

## Material and method

### 1. Fly Stocks

All *Drosophila melanogaster* lines were maintained on standard cornmeal–yeast–agar medium at 25 °C and 60% relative humidity, under a 12 h:12 h light–dark (LD) cycle, with a light intensity of approximately 500 lux during the light phase, unless otherwise specified. For the initial genetic screen, more than 50 RNAi lines targeting ubiquitin regulators were obtained from the Tsinghua Fly Center and the Vienna *Drosophila* Resource Center (VDRC). Full genotype information for all lines used in the screen is provided in Supplementary Table S1. Besides this following the other fly stocks used in this study: *TD2* (*tim-GAL4 ;UAS-dicer2*), *PD2* (*Pdf-GAL4 ;UAS-dicer2*), *TG4* (*tim-GAL4*), *TD2; Pdf-GAL80*, *Elav-GAL4* were all the GAL4 driver lines, *w^1118^* or *yw* as genetic background controls, the UAS-RNAi and overexpression lines including *Yippee*^RNAi-1^(VDRC46977), *Yippee*^RNAi-2^(VDRC108836), *Yippee^OE^-3×HA*, *CD8-GFP*(BL5137), *GFP^RNAi^*(BL44415), *per^RNAi^*(NIG2647R-2). UAS-RNAi and overexpression lines were all obtained from Bloomington *Drosophila* Stock Center (BDSC), the Vienna *Drosophila* Resource Center (VDRC) and the National Institute of Genetics Fly stock center (NIG-Fly).

### 2. *Drosophila* Behavioral Assays

Locomotor activity rhythms were monitored using the *Drosophila* Activity Monitoring System (TriKinetics, Waltham, MA). Male flies were collected within 3–6 days after eclosion and individual flies were then placed in glass tubes containing 5% sucrose and 2% agar as food entrained under a 12:12 h light-dark (LD) cycle at 25 °C and 60% humidity for 4 days, then kept under constant darkness (DD) for at least 6 days to assess free-running circadian rhythms. Activity data were collected in 1-min bins and analyzed using FaasX software and average activity actograms were generated using MATLAB software with the Sleep and Circadian Analysis Matlab Program (SCAMP) toolbox. Flies showing arrhythmic activity or mortality were excluded from analysis.

### 3. Immunofluorescence of *Drosophila* Brains

For both whole-brain staining and single-time-point sLNvs immunofluorescence, flies were entrained under standard LD cycles for three full days, and brains were collected at Zeitgeber Time 1 (ZT1). Flies were initially fixed in 4% paraformaldehyde (PFA) at room temperature for 1 hour. Adult fly brains were then dissected in PBST (0.1% Triton X-100 in PBS), followed by an additional fixation step in 4% PFA for 30 minutes. After fixation, brains were quickly rinsed three times in PBST, then washed slowly three times for 20 minutes each on a rotator. Samples were then blocked in 10% goat serum (prepared in PBST) for 1.5 hours at room temperature on a rotator. Brains were incubated overnight at 4 °C with the following primary antibodies diluted in 10% goat serum: rabbit anti-PER (1:1500, a gift from Joanna Chiu) and mouse anti-PDF (1:400, DSHB, AB360750). The following day, samples were washed six times in PBST for 20 minutes each on a rotator and incubated overnight at 4 °C in the dark with secondary antibodies: goat anti-mouse Alexa Fluor 488 (1:400) and Cy™5 AffiniPure Donkey Anti-Rabbit IgG (H+L) (1:400), both purchased from Jackson ImmunoResearch and diluted in PBST. Following secondary incubation, samples were washed six times in PBST (20 minutes each), then immersed in 70% glycerol for 30 minutes. Finally, brains were mounted and imaged using a Leica SP8 confocal microscope equipped with a 20× objective.

For multiple time point staining, flies were entrained under LD conditions for three full cycles and then transferred to constant darkness (DD), with samples collected at CT1 on the fourth day in DD. All other processing steps were identical to the single time point protocol.

### 4. RNA Extraction and Quantitative PCR (qPCR)

For *Yippee* knockdown efficiency detection, flies were entrained under LD conditions for three full cycles, and heads were collected at ZT1. For assessing the effect of *Yippee* downregulation on the expression of core clock genes, flies were entrained for three LD cycles and then transferred to constant darkness (DD); samples were collected at CT1 on the first day of DD. Flies were flash-frozen in liquid nitrogen, and heads were separated from bodies by vigorous vortexing. Total RNA was extracted using TRIzol reagent (Invitrogen) according to the manufacturer’s instructions. RNA concentration and purity were assessed using a NanoDrop spectrophotometer (Thermo Fisher Scientific). One microgram of total RNA was reverse-transcribed using the HiScript® III RT SuperMix for qPCR (+gDNA wiper) (Vazyme). Quantitative real-time PCR was performed using ChamQ Universal SYBR qPCR Master Mix (Vazyme) on a QuantStudio 6 Flex Real-Time PCR System (Thermo Fisher).

The following primers were used for qPCR:

***per***: Forward 5′-GACTCGGCCTACTCGAACAG-3′,

Reverse 5′-CGCGACTTATCCTTGTTGCG-3′

***tim***: Forward 5′-TGGACTGGTTACTAGCAACTCC-3′,

Reverse 5′-GGTCCTCATAGGTGAGCTTGT-3′

***clk***: Forward 5′-AATCGAGCCCAGACAGACG-3′,

Reverse 5′-GCATCCACTTTCTCCATCCCG-3′

***Yippee***: Forward 5′-TCTTCAATTGCGCCCAATGCCAC-3′,

Reverse 5′-ATGTGGCGACCCGTGAGCATGA-3′

***rpl32***: Forward 5′-CCGCTTCAAGGGACAGTATC-3′,

Reverse 5′-ACGTTGTGCACCAGGAACTT-3′

### 5. Western Blotting (WB) from Fly Heads

Flies were entrained under standard LD (12 h:12 h light–dark) conditions for three full cycles and collected at ZT1. Flies were flash-frozen in liquid nitrogen, and heads were separated from bodies by vortexing. Fly heads were homogenized in RIPA buffer (Beyotime) supplemented with protease and phosphatase inhibitor cocktails (Selleck), and lysates were incubated on 4℃ rotator for 30 minutes. Following lysis, samples were centrifuged at 12,000 rpm for 15 minutes at 4°C to remove debris. The supernatant was collected, mixed with 5× SDS–PAGE loading buffer (Beyotime), and boiled at 99°C for 10 minutes.

Equal amounts of protein (based on head number) were separated by SDS–PAGE and transferred to PVDF membranes (Millipore). Membranes were blocked with 5% non-fat dry milk in TBST (TBS with 0.1% Tween-20) for 1 hour at room temperature, followed by overnight incubation at 4°C with primary antibodies. The following primary antibodies were used: anti-HA (1:1000, mouse, Sigma, H3663) and anti-β-Actin (1:1000, mouse, Proteintech, 66009-1-Ig). After three washes with TBST, membranes were incubated with HRP-conjugated secondary antibodies (anti-mouse: 1:10,000, Jackson ImmunoResearch, 115-035-003; anti-rabbit: 1:10,000, Jackson ImmunoResearch, 711-035-152) for 1 hour at room temperature. Following another three washes with TBST, signals were detected using the SuperFemto ECL Chemiluminescence Kit (Vazyme) and visualized using the ChemiScope 6100 Touch Imaging System (Clinex).

### 6. Co-immunoprecipitation (CoIP) from Fly Heads

Flies were entrained under 12:12 LD conditions for three full cycles and collected at ZT1. After flash-freezing in liquid nitrogen, fly heads were separated from bodies by vortexing. Approximately 800 heads were homogenized in ice-cold IP lysis buffer (50 mM Tris-HCl, 150 mM NaCl, 1% NP-40, 1 mM EDTA, pH 7.5), supplemented with protease and phosphatase inhibitor cocktails (Selleck). The lysates were rotated at 4 °C for 30 minutes to ensure complete tissue lysis, followed by centrifugation at 12,000 rpm for 15 minutes at 4 °C. The supernatant was collected and a small portion was set aside as the input control. Input samples were mixed with 5× SDS–PAGE loading buffer (Beyotime), boiled at 99 °C for 10 minutes, and stored for western blot analysis.

The remaining lysate was precleared with 5 µL of pre-washed Dynabeads^TM^ Protein A/G(Invitrogen) (washed three times with cold lysis buffer, 5 minutes per wash on a rotator at 4 °C) for 30 minutes at 4 °C. Beads were removed, and the supernatant was divided equally into two tubes. One was incubated with a rabbit IgG isotype control antibody, and the other with an anti-PER antibody (rabbit-derived, a gift from Joanna Chiu). Samples were rotated overnight at 4 °C. On the following day, 10 µL of pre-washed beads were added to each sample and incubated for 3–5 hours at 4 °C with rotation to capture immune complexes. Beads were then washed three times with ice-cold lysis buffer (10 minutes per wash at 4 °C on a rotator). After the final wash, beads were resuspended in 60 µL of lysis buffer mixed with 5× SDS-PAGE loading buffer and boiled at 99 °C for 10 minutes to denature proteins and release them from the beads. The resulting samples were subjected to SDS-PAGE and analyzed by western blotting.

### 7. Cell Culture and Transfection

HEK293T cells were cultured in DMEM supplemented with 10% fetal bovine serum (FBS) and 1% penicillin–streptomycin (all from Gibco) at 37 °C in a humidified incubator with 5% CO₂. For both plasmid overexpression and *siRNA* knockdown experiments, HEK293T cells were transfected using Lipofectamine 3000 (Invitrogen) according to the manufacturer’s instructions. *siRNA* targeting *YPEL5* (*siYPEL5*) and negative control *siRNA* (all from GenePharma) were used at a final concentration of 50 nM. All downstream assays were performed 48–72 hours after transfection.

PER2::LUC U2OS cells were cultured in DMEM supplemented with 10% fetal bovine serum (FBS) and 1% penicillin–streptomycin (PS) at 37 °C in a humidified atmosphere containing 5% CO₂. For transfection, cells were seeded in 35-mm dishes and grown to 70%–90% confluence. Cells were then transfected with either *siYPEL5* or negative control *siRNA* using Lipofectamine 3000, following the manufacturer’s instructions.

### 8. Western Blotting in HEK293T Cells

HEK293T cells were transfected with the indicated plasmids or *siRNAs* and harvested 48–72 hours post-transfection. Cells were washed with ice-cold PBS and lysed in RIPA buffer (Beyotime) supplemented with protease and phosphatase inhibitor cocktails (Selleck). Lysates were incubated on ice for 30 minutes and centrifuged at 12,000 rpm for 15 minutes at 4 °C. The supernatant was collected, mixed with 5× SDS-PAGE loading buffer, and boiled at 99 °C for 10 minutes.

Equal amounts of protein were resolved by SDS–PAGE and transferred to PVDF membranes (Millipore). Membranes were blocked with 5% non-fat milk in TBST (TBS with 0.1% Tween-20) for 1 hour at room temperature, then incubated overnight at 4 °C with the following primary antibodies: anti-FLAG (1:1000, Sigma, F1804), anti-HA (1:1000, Sigma, H3663), anti-YPEL5 (1:1000, Proteintech, 11730-1-AP), anti-HSP90(1:1000, Proteintech, 13171-1-AP), and anti-β-actin (1:1000, Proteintech, 66009-1-Ig). After washing, membranes were incubated with HRP-conjugated secondary antibodies (Jackson ImmunoResearch, 1:10,000) for 1 hour at room temperature. Signals were detected using the SuperFemto ECL Kit (Vazyme) and visualized with the ChemiScope 6100 Touch Imaging System (Clinex).

### 9. Co-immunoprecipitation (CoIP) in HEK293T Cells

HEK293T cells were seeded in 10-cm dishes and transfected with different combinations of tagged expression plasmids using Lipofectamine 3000 (Invitrogen). For the PER2–YPEL5 interaction, cells were transfected with either PER2-FLAG or YPEL5-HA plasmids alone (single-transfection controls), or co-transfected with both constructs. After 48 hours, cells were lysed and immunoprecipitation was performed to assess specific interactions among the three groups.

In all CoIP experiments involving PER2, cells were treated with 10 μM MG132 (MCE) for 12 hours prior to harvest to prevent proteasomal degradation and improve detection of PER2-associated complexes. Cells were washed with ice-cold PBS and lysed in IP lysis buffer (50 mM Tris-HCl, pH 7.5, 150 mM NaCl, 1% NP-40, 1 mM EDTA) supplemented with protease and phosphatase inhibitor cocktails (Selleck). Lysates were incubated on ice for 30 minutes and centrifuged at 12,000 rpm for 15 minutes at 4 °C. The resulting supernatants were precleared with 5 μL of Dynabeads™ Protein A/G (Invitrogen) at 4 °C for 30 minutes.

After preclear, 1 μg of primary antibody (anti-FLAG, Sigma F1804) was added to the lysates and incubated overnight at 4 °C. The next day, 10 μL of Dynabeads™ Protein A/G were added for 3–5 hours at 4 °C to capture immune complexes. Beads were washed three times (10 minutes each) in cold IP buffer, resuspended in 60 µL of ice-cold IP lysis buffer, and mixed with 5× SDS–PAGE loading buffer. Samples were then boiled at 99 °C for 10 minutes to denature and release bound proteins. Input and immunoprecipitated samples were analyzed by western blotting using tag-specific primary antibodies to detect the presence of PER2, YPEL5.

### 10. Cycloheximide (CHX) Chase Assay

HEK293T cells were seeded in 10-cm dishes and transfected with a PER2-FLAG expression plasmid using Lipofectamine 3000 (Invitrogen). Six to eight hours later, cells were trypsinized and replated into 12-well plates at appropriate density. On the following day, cells were transfected with either *siYPEL5* or negative control *siRNA*. Forty-eight hours later, cells were treated with 100 μg/mL cycloheximide (CHX; MCE) to inhibit new protein synthesis. Samples were collected at 0, 4, 8, and 12 hours after CHX treatment for protein extraction and Western blot analysis of PER2-FLAG levels. To investigate the degradation pathway of PER2, HEK293T cells were transfected with the PER2-FLAG plasmid and replated into 12-well plates 6–8 hours post-transfection. After 48 hours, cells were divided into three groups and treated with 100 μg/mL CHX in combination with either vehicle (DMSO), 10 μM MG132 (MCE), or 10 μM chloroquine (CQ; Sigma-Aldrich, C6628). Protein samples were collected at 0, 4, 8, and 12 hours after treatment and subjected to Western blotting.

For all CHX assays, cells were washed with cold PBS and lysed in RIPA buffer (Beyotime) containing protease and phosphatase inhibitors (Selleck). Protein concentrations were normalized, and equal amounts of protein were separated by SDS–PAGE and analyzed by immunoblotting with anti-FLAG (1:1000, Sigma, F1804) and appropriate control antibodies as described above.

### 11. Ubiquitination Assay

To determine the type of ubiquitin linkage involved in YPEL5-mediated PER2 modification, HEK293T cells were co-transfected with PER2-FLAG and either wild-type ubiquitin, K48R mutant ubiquitin (lysine 48 mutated to arginine), or K63R mutant ubiquitin (lysine 63 mutated to arginine). Six to eight hours after the initial transfection, cells were replated into two 10-cm dishes. On the following day, cells were further transfected with either YPEL5-HA or *siYPEL5*, along with their respective negative controls. Forty-eight hours later, cells were treated with 10 μM MG132 (MCE) for 12 hours to inhibit proteasomal degradation and facilitate accumulation of ubiquitinated proteins. Cells were harvested the next day.

Sample preparation, immunoprecipitation, and Western blotting were performed as described in the CoIP section. PER2-FLAG was immunoprecipitated, and ubiquitination levels were assessed by probing the immunoblots with an anti-HA antibody (1:1000, Sigma, H3663), which specifically detects HA-tagged ubiquitin conjugated to PER2.

### 12. Immunoprecipitation–Mass Spectrometry (IP-MS)

Immunoprecipitation followed by mass spectrometry (IP-MS) was performed on head extracts from *TD2/+* and *TD2/+; Yippee^OE^-HA/+* flies collected at ZT1, following three full cycles of LD entrainment. Approximately 1,200 adult fly heads per sample were flash-frozen in liquid nitrogen, vortexed to separate the heads, and homogenized in ice-cold IP lysis buffer (50 mM Tris-HCl pH 7.5, 150 mM NaCl, 1% NP-40, 1 mM EDTA) supplemented with protease and phosphatase inhibitors (Selleck). Lysates were incubated on ice for 30 minutes and clarified by centrifugation at 12,000 rpm for 15 minutes at 4 °C. Supernatants were precleared with 5 μL Dynabeads™ Protein A/G (Invitrogen) for 30 minutes at 4 °C with rotation. After removal of the beads, 1 μg of anti-HA antibody (Sigma, H3663) was added to each sample and incubated overnight at 4 °C. The following day, 10 μL Dynabeads™ Protein A/G were added for an additional 3-5 hours of incubation at 4 °C. Beads were washed three times with cold lysis buffer (10 minutes per wash), and bound proteins were eluted using 8 M urea solution to disrupt protein–bead interactions.

The urea-eluted proteins were submitted to the laboratory of Professor Pan Fang at the Institute of Molecular Enzymology, Soochow University, for sample preparation. Briefly, 25 μL of each eluate was mixed with 75 μL of 50 mM HEPES, followed by reduction and alkylation with TCEP and CAA, respectively^[36, 37]^. The urea concentration was then further diluted, and the samples were digested with trypsin, desalted, and quantified^[38]^. Finally, 5 μg peptides from each sample were submitted by Professor Fang’s group to an external service provider for LC–MS/MS analysis.

Mass spectrometry data were searched against the *Drosophila melanogaster* UniProt database (including both reviewed and unreviewed entries). Processed results were returned to us for further bioinformatic analysis. Protein–protein interaction networks were analyzed using STRING (https://string-db.org/). Gene Ontology (GO) and KEGG pathway enrichment analyses were performed using DAVID Bioinformatics Resources. Structural predictions of selected proteins and their interactions were conducted using AlphaFold3.

### 13. Circadian Bioluminescence Recording in U2OS Cells

PER2::LUC U2OS cells were transfected with *siYPEL5* or negative control *siRNA* using Lipofectamine™ 3000 (Invitrogen) as described above. Forty-eight hours post-transfection, the culture medium was replaced with D2902 recording medium to synchronize cellular circadian oscillations and enable bioluminescence recording.

D2902 medium was prepared by dissolving 10 g of D2902 powder (Sigma-Aldrich, Lot No. 0000349350) in water to a final volume of 1 L, containing 7.5% NaHCO₃ (4.7 mL), 3.5 g glucose, and 10 mL of 1 M HEPES, with pH adjusted to 7.2. Before use, the medium was supplemented with 10% fetal bovine serum (FBS), 1% penicillin–streptomycin (PS), 0.1 μM dexamethasone, and 0.1 mM D-luciferin (GoldBio, CAS No. 115144-35-9). Each culture dish was sealed with optically clear qPCR sealing film and placed into a LumiCycle luminometer maintained at 37 °C. Bioluminescence was recorded continuously for approximately 7 days.

### 14. Mouse SCN Injection, Wheel-Running Behavior, and Immunostaining

#### 14.1 Animals and Stereotaxic Injection

Eight-week-old male *Ypel5^Flox/Flox^* mice were used for bilateral stereotaxic injections targeting the suprachiasmatic nucleus (SCN). Mice were anesthetized with 2.5% isoflurane for induction and maintained at 1.5% during the procedure. The head was secured in a stereotaxic apparatus, and the scalp was shaved and sterilized with 75% ethanol. After a midline incision, the skull was cleaned with 3% hydrogen peroxide to visualize the cranial sutures.

Glass electrodes were used to mark the bregma (X=0, Y=0, Z=0), and head leveling was performed by comparing Z-axis readings 1 mm lateral to bregma (left and right), adjusting until the difference was within ±0.01 mm. Anterior–posterior alignment was confirmed by moving 4.2 mm posterior to identify lambda, ensuring the Z-axis difference from bregma was within ±0.02 mm. A small burr hole (∼1.5 mm diameter) was drilled at the target coordinates, and the dura was carefully removed. Skull thickness was measured and subtracted from the target depth (Z=–5.85 mm) to determine the precise injection depth. Viral vectors VGAT1-eGFP (control group) and VGAT1-Cre (experimental group) were used. A total of 150 nL of virus was delivered into each SCN hemisphere at a rate of 100 nL/min. The injection pipette was held in place for 10 minutes post-injection before being slowly withdrawn. The contralateral hemisphere was injected using the same procedure. Following injection, the skull was dried and the incision was sutured. Mice were returned to individual cages for recovery.

#### 14.2 Wheel-Running Activity Monitoring

Three weeks post-injection, *Ypel5^Flox/Flox^* mice were individually housed in cages equipped with running wheels. Mice were first entrained under a 12 h:12 h light–dark (LD) cycle for 7–14 days, then transferred to constant darkness (DD) for at least 14 days to assess endogenous circadian rhythms.

Wheel-running activity was continuously monitored using the ClockLab data acquisition system. Circadian period was analyzed using ClockLab and GraphPad Prism. Mice had free access to food and water throughout the entire recording period.

#### 14.3 Immunofluorescence Validation of Injection Site

To verify the accuracy of viral injection into the SCN, mice were deeply anesthetized and transcardially perfused with 4% paraformaldehyde (PFA). Brains were post-fixed overnight in 4% PFA at 4 °C, then coronally sectioned at 40 μm thickness. Brain slices were mounted and counterstained with DAPI for 5 minutes in the dark to label nuclei. EGFP fluorescence was examined using an OLYMPUS IX53 microscope (10× objective) to confirm viral expression within the SCN region.

Mouse stereotaxic injections and behavioral recordings were carried out with support from the laboratory of Professor Ying Xu at the Cambridge–Soochow University Genomic Resource Center, Soochow University.

### 15. Statistical Analysis

All statistical analyses were performed using GraphPad Prism (version 8.0.1) and R (version 4.3.2). Data are presented as mean ± SEM unless otherwise indicated.

For pairwise comparisons between two groups, unpaired two-tailed Student’s *t*-tests were used. For comparisons involving more than two groups, one-way or two-way analysis of variance (ANOVA).

For *Drosophila* behavioral assays, period length and rhythmic power were calculated from raw activity data using GraphPad Prism. Representative actograms were generated using a custom MATLAB script.

For qPCR data, relative gene expression was calculated using the ΔΔCt method. Western blot and fly brain immunostaining intensities were quantified using ImageJ, and group differences were assessed using *t*-tests or ANOVA as appropriate.

For CHX assays, PER2 protein band intensities at each time point were normalized to time zero, and degradation curves were plotted using GraphPad Prism. Group comparisons across time points were performed using two-way ANOVA.

For mouse wheel-running data, circadian period was determined using ClockLab-based analysis, and subsequent statistical comparisons were performed in GraphPad Prism.

For U2OS PER2-luc recordings, circadian period and amplitude were extracted using LumiCycle Analysis software. Statistical comparisons of period and amplitude between groups were performed using unpaired *t*-tests.

A *p*-value < 0.05 was considered statistically significant. Significance was indicated as *P* < 0.05 (*), *P < 0.01* (**), *P < 0.001* (***), and *P* < 0.0001 (****).

## Acknowledgments

We thank Dr. Ying Xu and her lab (Soochow University) for providing technical assistance with stereotaxic injections and behavioral recordings in mice. We are especially grateful to Yizhun Zeng for his invaluable help with surgical procedures and animal handling. We also thank Dr. Joanna Chiu (University of California, Davis) for generously providing the anti-PER antibody used in *Drosophila* experiments. We acknowledge the laboratory of Dr. Pan Fang (Institute of Molecular Enzymology, Soochow University) for assistance with sample preparation and proteomic analysis of IP–MS data. We sincerely thank Dr. Guoqiang Xu’s laboratory (Soochow University) for their insightful guidance on the design and execution of ubiquitination-related experiments. This study was supported by National Natural Science Foundation of China (32271206 to Y.Z.) and Postgraduate Research & Practice Innovation Program of Jiangsu Province (KYCX25_3478 to M.L.).

---

**Fig S1.**
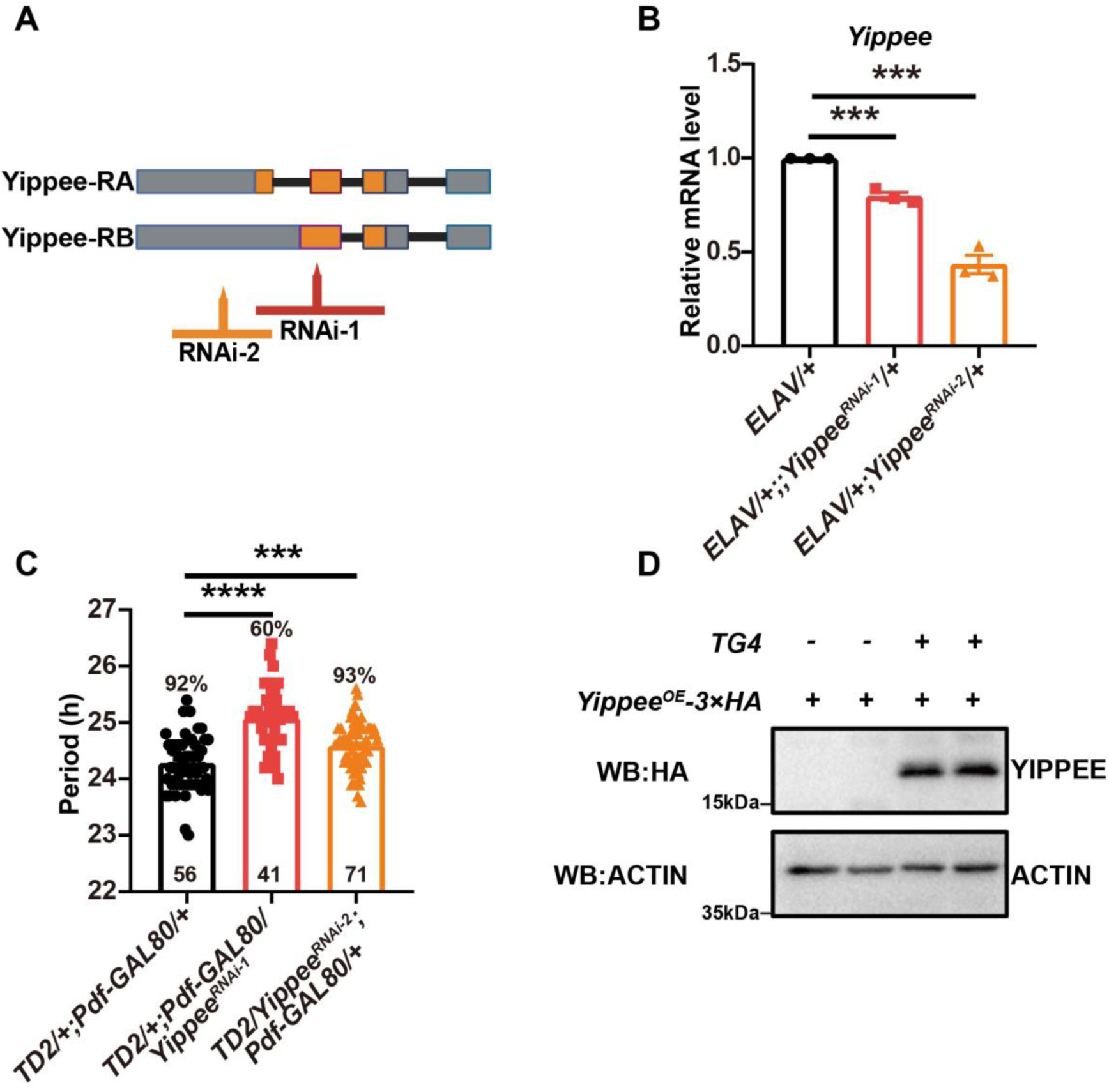
*Yippee* knockdown efficiency and its primary neuronal targets. (A) Schematic diagram showing the target sites of the RNA interference in two independent *Yippee* RNAi lines. RNAi-1 and RNAi-2 correspond to VDRC46977 and VDRC108836, respectively. (B) Knockdown efficiency of the two *Yippee* RNAi lines using the pan-neuronal driver *ELAV-GAL4*. Statistical analysis was performed using an unpaired t-test. Error bars indicate SEM. ***: *P* < 0.001. (C) Influence of *Yippee* knockdown on circadian rhythms in PDF negative neurons. Statistical analysis was performed using one-way ANOVA. Error bars represent SEM. ***: *P* < 0.001, ****: *P* < 0.0001. (D) Western blot analysis confirming *Yippee* overexpression. *TG4* denotes *tim-GAL4*; *Yippee^OE^* denotes *UAS-Yippee^OE^-3×HA*. Anti-HA antibody was used to detect YIPPEE level. Actin served as the internal loading control.

**Fig S2.**
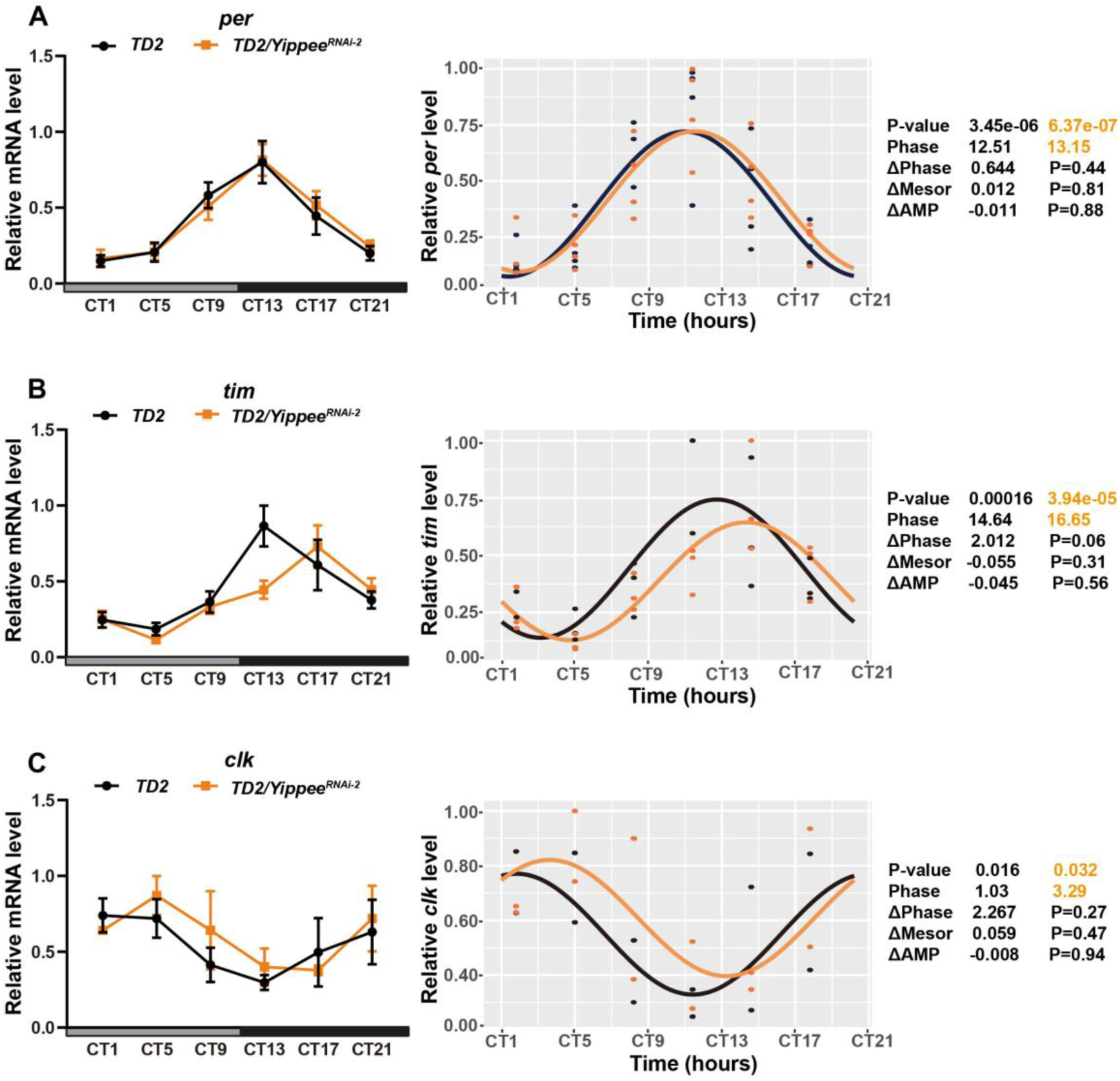
Downregulation of *Yippee* does not affect the mRNA expression of core clock genes. (A–C) RT-qPCR analysis of *per*, *tim*, and *clock* mRNA levels in *Yippee* RNAi flies. Flies were entrained under 12:12 h light–dark (LD) conditions for three days, then transferred to constant darkness (DD). Samples were collected every four hours over a 24-hour period in DD. Rhythmic expression profiles were analyzed using Circacompare in R Studio. Right panels show rhythm parameters; *P* > 0.05 indicates no significant difference between groups.

**Fig S3.**
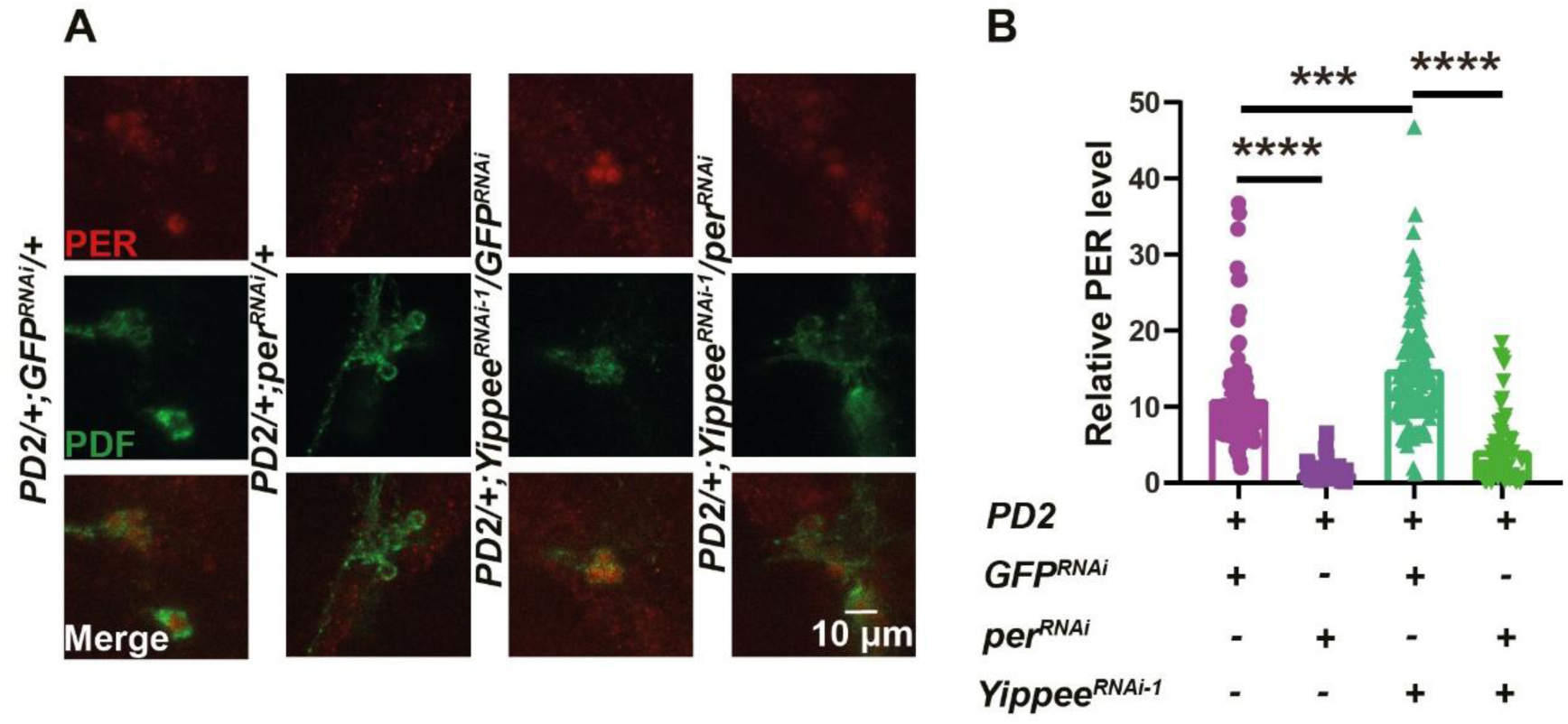
Verification of PER levels in *per* and *Yippee* double knockdown flies. (A) Double knockdown of *per* and *Yippee* partially reverses the elevated PER levels observed in *Yippee* knockdown flies. Representative confocal images showing PER (red) and PDF (green) immunostaining in small ventrolateral neurons (sLNvs) across four genotypes: *PD2/+;GFP^RNAi^/+*; *PD2/+;per^RNAi^/+*; *PD2/+; Yippee^RNAi-1^ /GFP^RNAi^*; *PD2/+; Yippee^RNAi-1^ /per^RNAi^*. Flies were entrained under a 12:12 light–dark (LD) cycle for three days and collected at Zeitgeber Time 1 (ZT1). PDF immunostaining was used to identify the sLNvs. (B) Quantification of PER fluorescence intensity in sLNvs across the four groups. Data are presented as mean ± SEM. Statistical significance was determined by unpaired *t*-tests. ***: *P* < 0.001; ****: *P* < 0.0001.

**Fig S4.**
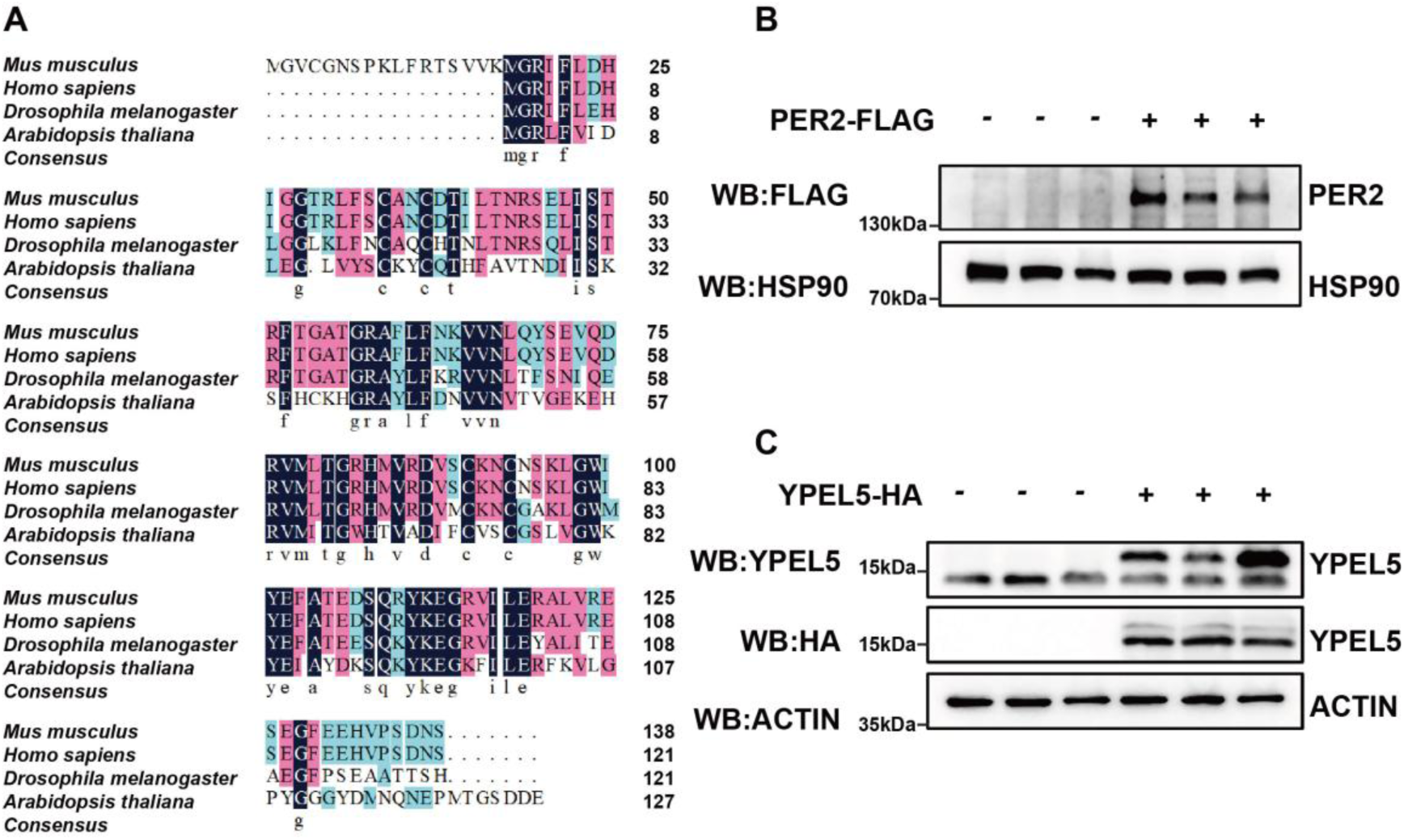
YIPPEE protein is evolutionarily conserved across species. (A) YIPPEE protein sequence alignment across species. Sequence alignment was performed using DNAMAN to compare YIPPEE protein sequences from *Mus musculus*, *Homo sapiens*, *Drosophila melanogaster*, and *Arabidopsis thaliana*. Dark blue indicates highly conserved residues; pink or light blue indicate moderately conserved residues; white or unshaded regions represent non-conserved sites. (B) Western blot analysis confirming PER2-FLAG expression in HEK293T cells. Anti-FLAG antibody was used to detect PER2. HSP90 served as the internal loading control. (C) Western blot analysis confirming YPEL5-HA expression in HEK293T cells. Anti-HA and anti-YPEL5 antibodies were used to detect YPEL5. HSP90 served as the internal loading control.

**Fig S5.**
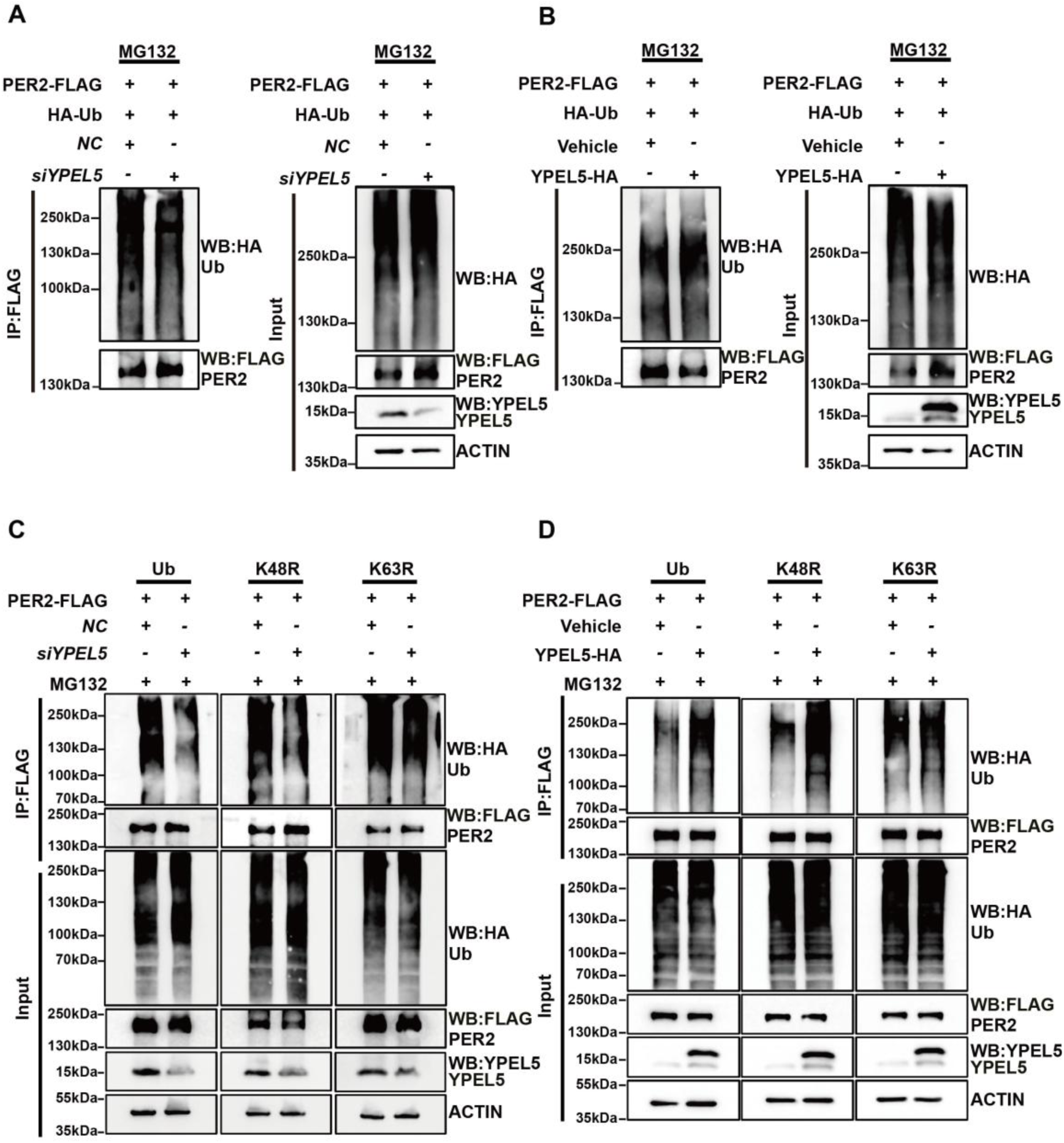
YPEL5 promotes the ubiquitination of PER2 through K63-linked modification. (A–B) YPEL5 modulates the ubiquitination of PER2. These are independent replicate experiments validating the findings shown in Fig. 4A and 4B.(C–D) YPEL5 regulates the K63-linked ubiquitination of PER2. These panels present repeated experiments corresponding to Fig. 4C and 4D.

**Fig S6.**
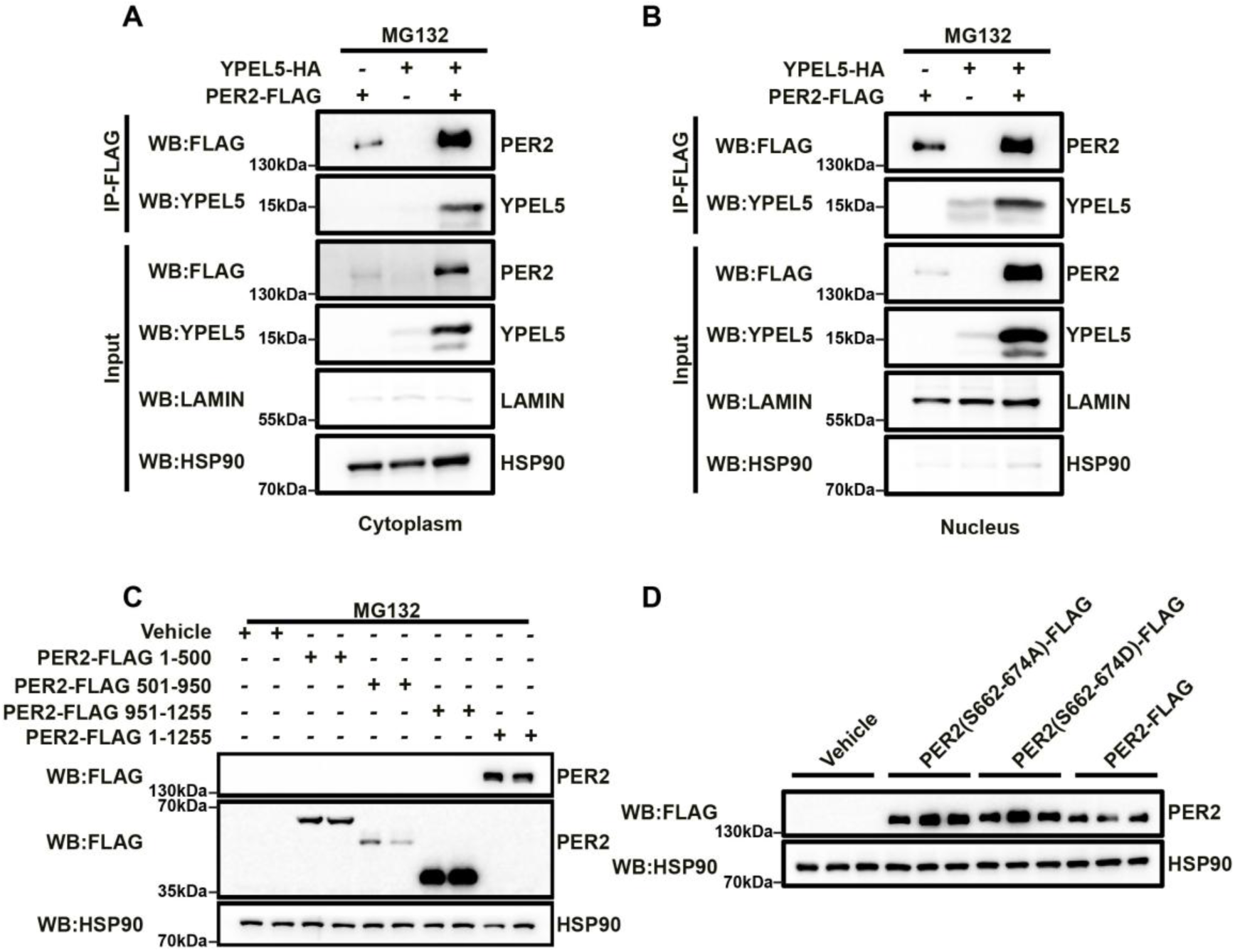
Expression of PER2 and related constructs used in phosphorylation assays. (A-B) Co-immunoprecipitation analysis of YPEL5 and PER2 interactions in cytoplasmic and nuclear fractions. Panel A shows the interaction between YPEL5 and PER2 in the cytoplasmic fraction, while Panel B shows their interaction in the nuclear fraction. HSP90 and Lamin were used as cytoplasmic and nuclear markers, respectively, to verify fractionation efficiency. (C) Western blot analysis confirming the expression of PER2 truncation constructs corresponding. The fragments 1–500 aa, 501–950 aa, and 951–1255 aa represent distinct regions of PER2. (D) Western blot analysis validating the expression of PER2 phosphorylation mutant constructs. PER2(S662–674A) denotes alanine substitution of residues S662, S665, S668, S671, and S674, whereas PER2(S662–674D) denotes aspartic acid substitution at these sites.

**Table S1.**
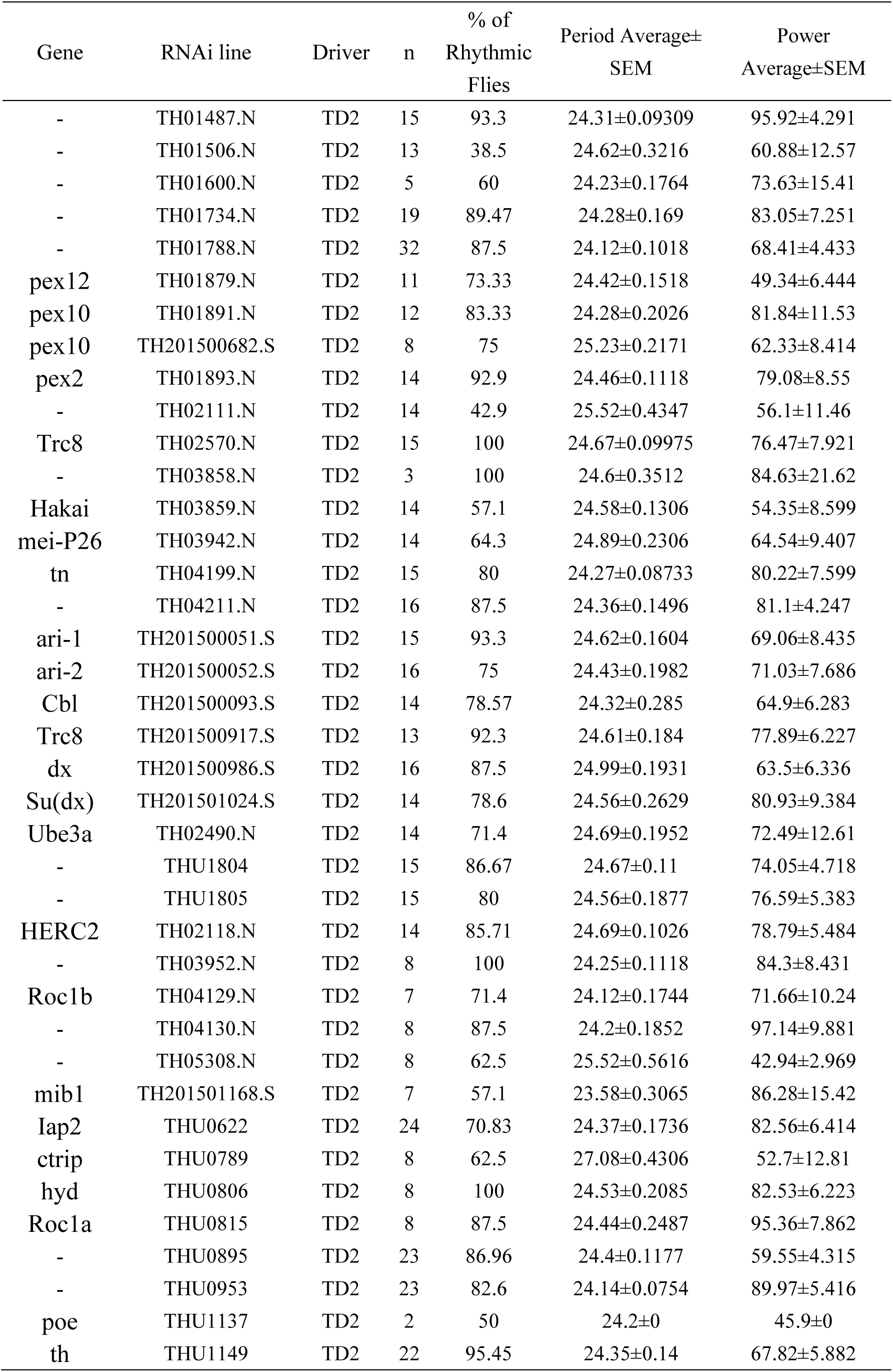
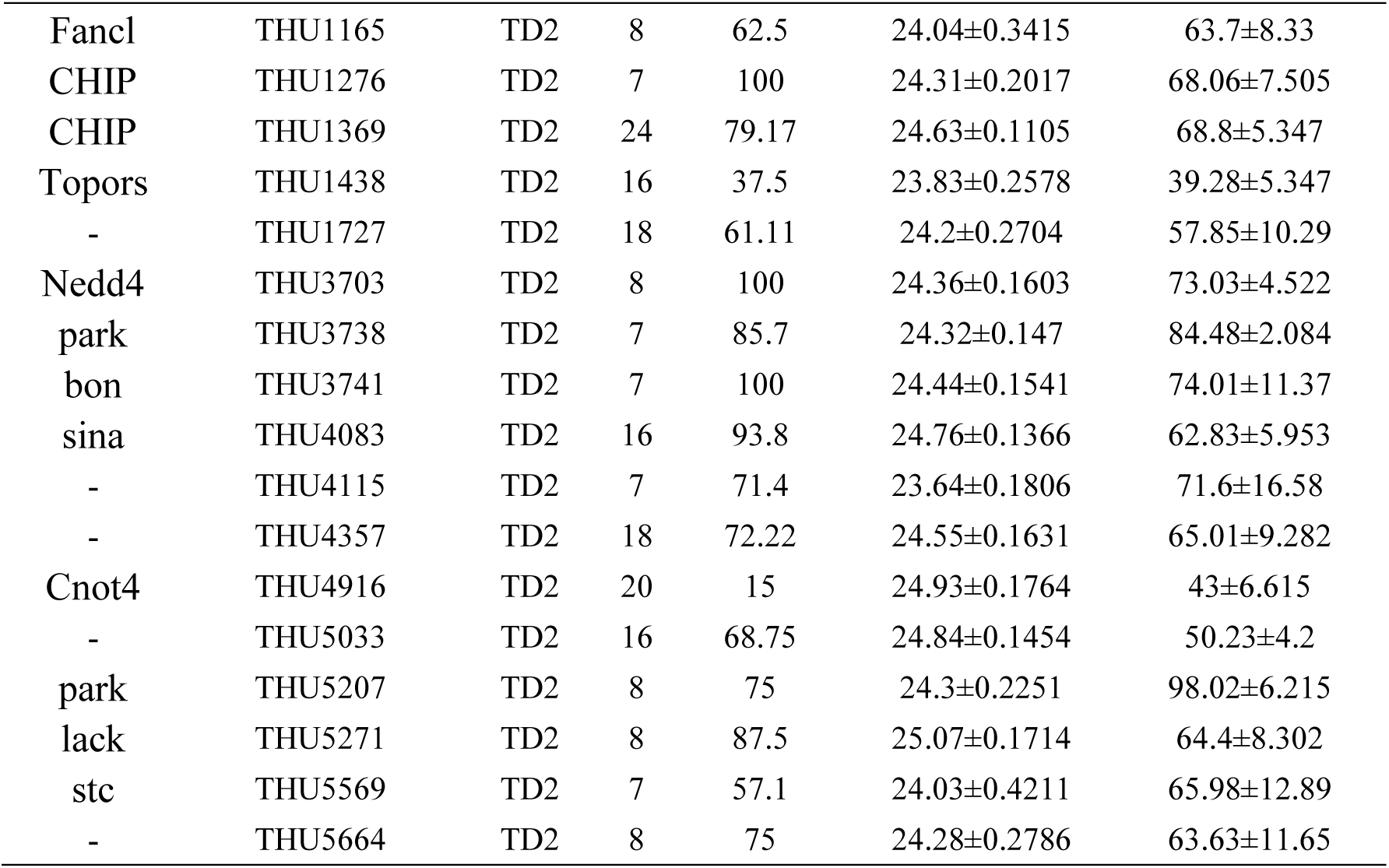
Circadian behaviors of ubiquitin-related genes knockdown.

